# Multichannel optical cochlear implants enable spectrally distinct auditory activity

**DOI:** 10.64898/2026.05.15.725096

**Authors:** Niels Albrecht, Elisabeth Koert, Anna Vavakou, Lennart Roos, Lukasz Jablonski, Juan Pablo Marcoleta, Josep Cardona Audi, Jette Alfken, Eric Klein, Patrick Ruther, Thomas Mager, Kathrin Kusch, Tobias Moser

## Abstract

When hearing fails, cochlear implants (CIs) restore auditory perception. Yet, coding of spectral information remains a bottleneck because each electrode broadly activates the auditory nerve. As light can be more conveniently confined, optical (o)CIs present an alternative. Here, we combined expression of the potent channelrhodopsin ChReef in spiral ganglion neurons (SGNs) with oCIs comprising 5–10 green LEDs in gerbils. This combination enabled systematic comparison of encoding intensity and spectral information by individual oCI channels to acoustic and electrical stimulation within a translationally feasible energy range. Recordings from the inferior colliculus (ICC) showed that ChReef aligned SGN light sensitivity with LED radiant fluxes: ICC activity had thresholds <200 nJ and reached a maximum equivalent to that achieved with 51 dB SPL pure tones. Multichannel oCIs enabled tonotopically ordered, spectrally distinct stimulation that more closely resembled acoustic stimulation than did electrical stimulation. Some LEDs elicited multiple spectral peaks at higher intensities. Linear discriminant analysis of ICC activity indicated improved channel discriminability for optical over electrical stimulation. In summary, µJ-oCI-stimulation achieves improved spectral resolution over state-of-the-art electrical stimulation.

**The Paper Explained:** *Problem:* Electrical cochlear implants (eCIs) partially restore speech comprehension in most of >1 million otherwise deaf users, who still face challenges hearing in daily situations. This is primarily due to poor spectral selectivity of electrical sound encoding. Spatially more confined optogenetic activation of the auditory nerve by optical cochlear implants (oCI) promises to overcome this limitation. However, a thorough characterization of bionic coding of sound information by multichannel oCI in a clinically translatable energy range is needed to evaluate the potential for improved hearing restoration.

*Results:* Here, we combine the potent channelrhodopsin ChReef and 10-channel oCI based on green LEDs in gerbils and characterize their utility for encoding of spectral and intensity information by individual channels using multielectrode array recordings from the midbrain. ChReef enabled activation of the auditory pathway with nJ thresholds and midbrain activity equivalent to that achieved with 51 dB SPL pure tones with low µJ radiant energy. The cochlear spread of excitation and channel discriminability in ICC activity substantially narrowed the gap to acoustic stimulation compared to state-of-the-art electrical stimulation.

*Impact:* Our work demonstrates great potential of multichannel optogenetic stimulation for encoding sound frequency information within a translationally feasible energy range.

## Introduction

With a prevalence of 5.6 % disabling hearing loss (HL) is the most common human sensory deficit (WHO, 2021). Sensorineural HL, caused by dysfunction of the cochlea, is the most frequent form. While inner ear gene therapy is emerging and will serve select groups of patients with a given genetic HL (e.g. Lv et al., 2024), hearing aids and electrical cochlear implants (eCIs) remain key means of rehabilitation (Moser et al., 2024). CIs bypass the dysfunctional organ of Corti and directly stimulate spiral ganglion neurons (SGNs) (Wohlbauer and Dillier, 2025; Zeng, 2017). eCIs provide open speech perception for most users after a year of rehabilitation. Yet, eCI users typically face difficulties in speech comprehension in daily situations with noisy backgrounds or competing sound sources (Hunniford et al., 2023). This shortcoming is commonly related to wide current spread activating large populations of SGNs (Kral et al., 1998; Miller et al., 2006; Shannon, 1983). The spiral ganglion, like the sensory organ of Corti, shows a sound-frequency spatial (tonotopic) organization, offering “labeled lines” for sound frequencies to the brain. Nowadays, eCIs take limited advantage of this frequency map, although current focusing techniques can provide narrower spread of excitation (Bierer, 2010). Optical CIs (oCI) promise improved frequency coding, as light can be confined in space better than electrical current, and, consequently, better hearing restoration for CI users (Hernandez et al., 2014; Izzo et al., 2006). Optogenetic stimulation of SGNs utilizes channelrhodopsins (ChRs), light-gated ion channels (Hernandez et al., 2014; Mager et al., 2018; Wrobel et al., 2018). Indeed, optogenetic SGN stimulation showed improved spectral selectivity compared to eCI stimulation (Dieter et al., 2020, 2019; Keppeler et al., 2020).

However, the power requirement has remained a challenge for optogenetic SGN stimulation to become available for clinical hearing restoration (Huet et al., 2024). In fact, despite the use of the powerful channelrhodopsin (ChR) 2 variant CatCh (Kleinlogel et al., 2011), optical energy thresholds (∼2 µJ) for SGN activation were more than an order of magnitude larger than the electrical ones (∼0.05 µJ, (Zeng, 2017)). Note, that increased number of channels, losses of energy conversion and imperfect light coupling need to be taken into consideration when comparing power budgets of the oCI and eCI. This challenged the utility of implantable microscale emitters for preclinical characterization of multichannel oCI (Dieter et al., 2020; Keppeler et al., 2020). With previous µLED-based approaches, the light sensitivity of CatCh was frequently insufficient for activation by individual LEDs, requiring simultaneous operation of multiple LEDs (Dieter et al., 2020). Moreover, reliable responses from all channels implemented on multichannel oCI prototypes were not consistently observed within the same animal (Keppeler et al., 2020). Thus, achieving energy-efficient and tonotopic activation of the spiral ganglion by individual oCI channels remained an important objective for characterizing optogenetic stimulation by multichannel oCIs.

The recent engineering and cochlear application of ChReef, an optimized larger-conductance green light activated ChR, provided energy thresholds of 0.2 µJ and preliminary data indicated a better match of this to LED-based oCI (Alekseev et al., 2025). Although the temporal properties of ChReef (*τ*_off_ ∼30 ms at physiological temperature) challenge its use for clinical hearing restoration, the high light sensitivity it conveys to SGNs paved the way for preclinical characterization of the utility of multichannel oCI at a power budget more compatible with clinical translation. However, the spectral and intensity encoding enabled by the combination of LED-oCIs with ChReef had not yet been systematically characterized.

Here, we used AAV-based expression of ChReef in SGNs and stimulation by individual green LEDs to characterize preclinical multichannel oCI with recordings from the central nucleus of the inferior colliculus (ICC) in Mongolian gerbils. The ICC is an auditory midbrain hub representing most if not all information relevant to assessing sound encoding in the cochlea (Drotos and Roberts, 2024). Recordings of multiunit activity from multielectrode arrays inserted along the tonotopic axis of the ICC offers convenient access to estimate which frequency range of the cochlea is activated by natural sound, eCI or oCI stimulation (Dieter et al., 2019; Matic et al., 2011; Snyder et al., 2004). The gerbil offers a cochlear size amenable to insertion of preclinical oCI (Dieter et al., 2020; Keppeler et al., 2021, 2020).

Here, we demonstrate that ChReef-mediated oCI stimulation provides efficient and spatially confined activation of the auditory pathway. We consistently found responses to µJ - stimulation by individual channels implanted into the gerbil cochlea, overcoming previous limitations in reliable single-emitter activation. The maximal ICC activity elicited by single emitters reached that elicited by 51 dB SPL pure tones. Consistent with previous studies, we demonstrate that optogenetic stimulation substantially improved spectral selectivity compared with electrical stimulation but did not achieve that of acoustic stimulation. Going beyond previous work, reliable activation by individual emitters also enabled us to assess channel discriminability at the level of ICC activity. ICC responses elicited by different oCI channels were more discriminable than those elicited by eCI channels, particularly for stimulus intensities eliciting high activity levels. By taking advantage of postmortem micro-computed tomography, we related emitter position and orientation to the physiologically derived estimates of optogenetic SGN activation The tonotopic locations of ICC responses closely matched those expected from the intracochlear emitter positions based on the Greenwood function. The high light sensitivity conveyed by ChReef also revealed a thus far undetected additional peak of activation beyond the targeted tonotopic position, which we carefully characterized. In summary, the results demonstrate selective and efficient ChReef-mediated encoding of frequency and intensity information.

## Results

### Characterizing optogenetic cochlear stimulation by multichannel oCI using recordings from the auditory midbrain

To provide robust ChR expression in SGNs, we injected AAV-PHP.S carrying ChReef under control of the human synapsin promoter into the cochlea of postnatal Mongolian gerbils (p8-9). After a minimum of 8 weeks, we first assessed successful transduction using optically evoked auditory brainstem responses (oABRs) elicited by a single optical fiber placed at the round window. oABRs to sub-µJ light pulses were observed in 13 out of 15 gerbils (0.32 [0.24, 0.44] µJ (median [IQR: 25^th^,75^th^ percentile], n=13, **Fig. EV1**). Successful ChReef expression was confirmed by postmortem immunohistochemistry and confocal microscopy: 24 to 74 % of SGNs were ChReef-positive in animals with oABR thresholds < 10 µJ (**Fig. EV2B**). In these animals, we observed a ∼30% lower median SGN density in injected cochleae compared with both the non-injected contralateral cochleae and untreated control cochleae. This difference was significant compared with left control cochleae (p = 0.002) but not compared with the untreated contralateral cochleae of the same animals (p = 0.17; **Fig. EV2C**). The SGN density reduction was strongest in the apical turn of the cochlea (**Fig. EV2C**).

When oABRs were detected at energies below 10 µJ, we continued to assess spectral and intensity coding by ChReef-mediated optogenetic SGN stimulation using multichannel oCI assembled from up to ten LEDs integrated on polyimide substrates (LED-oCI, Keppeler et al., 2020). Green LEDs (527 nm) provide a good match to ChReef (absorption maximum: 520 nm, (Alekseev et al., 2025). We first placed a 32-channel linear recording array into the right ICC of ChReef-treated animals and a group of non-injected animals for acoustic stimulation (**Fig. 1A**). Complete ICC recordings were obtained from 9 of 13 animals with oABR thresholds <10 µJ (loss due to the long duration and complexity of the experiments) and from 8 non-injected control animals. In the acoustic control animals, poloxamer hydrogel was applied to the right ear canal and bulla to induce conductive hearing loss and thereby predominantly restrict acoustic stimulation to the left ear, matching the side used for oCI stimulation in injected animals. The ChReef-transduced animals were not deafened, allowing placement along the tonotopic axis of the ICC to be assessed using pure-tone stimulation across a range of frequencies and intensities in both groups. Linear fits of frequency versus best-responding (lowest-threshold) electrode position were used to derive characteristic frequencies (C_f_) and estimate tonotopic slopes (**Appendix Fig. S1**). We found that the median tonotopic ICC slope, expressed as the change in C_f_ relative to recording position along the ICC axis, amounted to 4.75 oct/mm which closely relates to literature values (Dieter et al., 2020, 2019; Harris et al., 1997; Schnupp et al., 2015).

**Figure 1.**
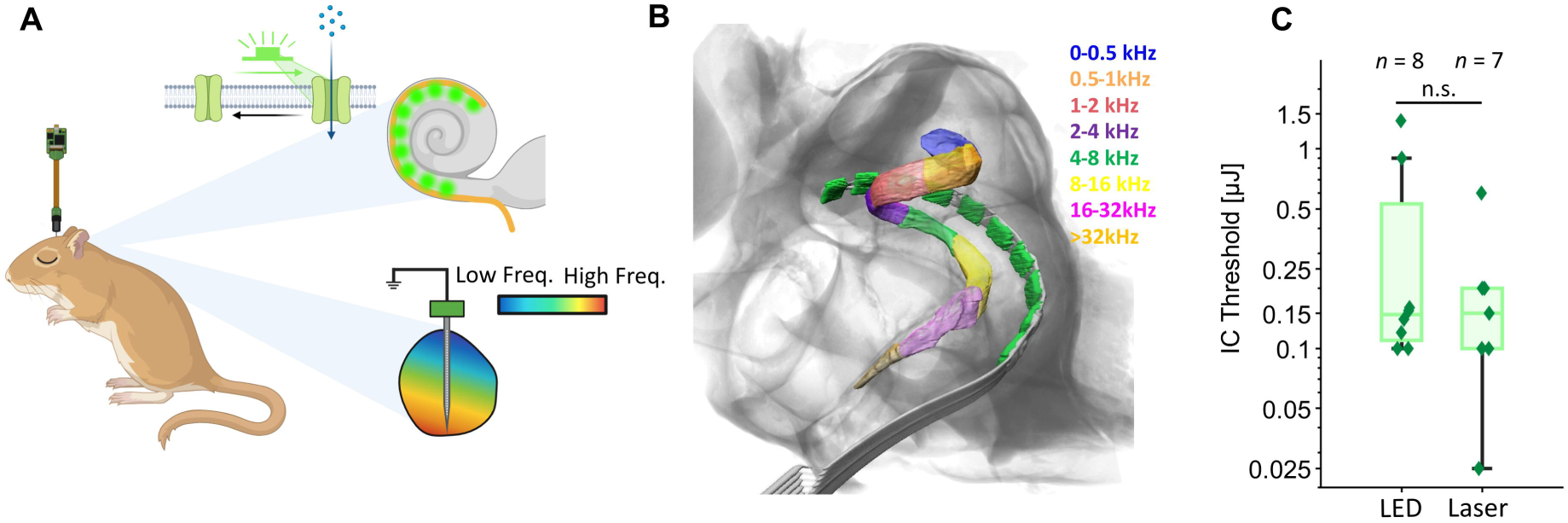
Experimental approach. **(A)** Optically evoked multi-unit activity was recorded at 32 recording sites along the tonotopic axis of the central nucleus of the inferior colliculus (ICC). Stimulation was delivered via optical cochlear implants (oCI) carrying up to 10 independently addressable LED emitters. Created in BioRender. Albrecht, N. (2026) https://BioRender.com/29xf30k. **(B)** X-ray tomography of a gerbil cochlea implanted with an oCI. The implant was fixed in the position used for stimulation for subsequent extraction and imaging of the cochlea. Frequency areas of the Rosenthal’s canal are highlighted according to the gerbil Greenwood function (Müller, 1996). Individual LEDs measured 220 × 270 µm (width × length along the implant). **(C)** Threshold of LED- and laser fiber-mediated ICC activation, defined as the first stimulation intensity eliciting a d’ ≥ 1. Thresholds across LEDs within the same animal were averaged before plotting (N = 8 gerbils for LED, N = 7 gerbils for laser fiber). Statistical significance was tested by Wilcoxon signed-rank test (n.s, non-significant; p > 0.05). Boxes indicate quartiles and median, with whiskers extending to the minimum and maximum values. Exact p-values are reported in **Table EV1**.

We then implanted LED-oCI with emitter spacings of either 300 or 400 µm via the round window or a cochleostomy in the basal turn of the cochlea (**Fig. 1 A, B**). Of the nine injected animals with successful ICC recordings, one was implanted through a mid-turn cochleostomy and was excluded to ensure comparability across animals. Of the eight included animals, three received a full 10-channel insertion and five received a 5-channel insertion; one additional 5-channel insertion was performed in one of the animals that also received a 10-channel insertion. In all implanted animals, 1 ms light pulses delivered from individual LEDs reliably evoked ICC activity with a median threshold of 0.15 [0.11, 0.53] µJ (n=8, **Fig. 1C**). We used such brief optical pulses (1 ms) of varying intensity from individual LEDs to stimulate the auditory system and compared the resulting ICC activity to that evoked by pure tones in non-injected animals. Moreover, we compared it to previously recorded ICC activity evoked by monopolar electrical stimulation in 12 wild-type gerbils (Dieter et al., 2019). To maximize comparability, the present recordings were obtained using the same experimental setup, and both datasets were processed using the same analysis pipeline.

We first computed evoked firing rates (evoked rate, ER) as a basic measure of activity level in defined time windows (**Fig. 2A-C).** Spontaneous firing rates measured before stimulation were subtracted from the firing rate within a 13-ms window following response onset to obtain evoked firing rates. This baseline correction was particularly important because spontaneous rates differed between modalities (**Appendix Fig. S2**). We note that interpretation of this comparison needs to consider the difference in stimulus duration. Compared to a recent study using acoustic clicks, i.e. with comparable duration (Michael et al., 2023), this study aimed to characterize spectral coding and, hence relied on pure tones. Therefore, to maintain comparability across stimuli of different durations, we calculated firing rates in a 13 ms window following the response latency, thereby capturing the onset response to each stimulus.

**Figure 2.**
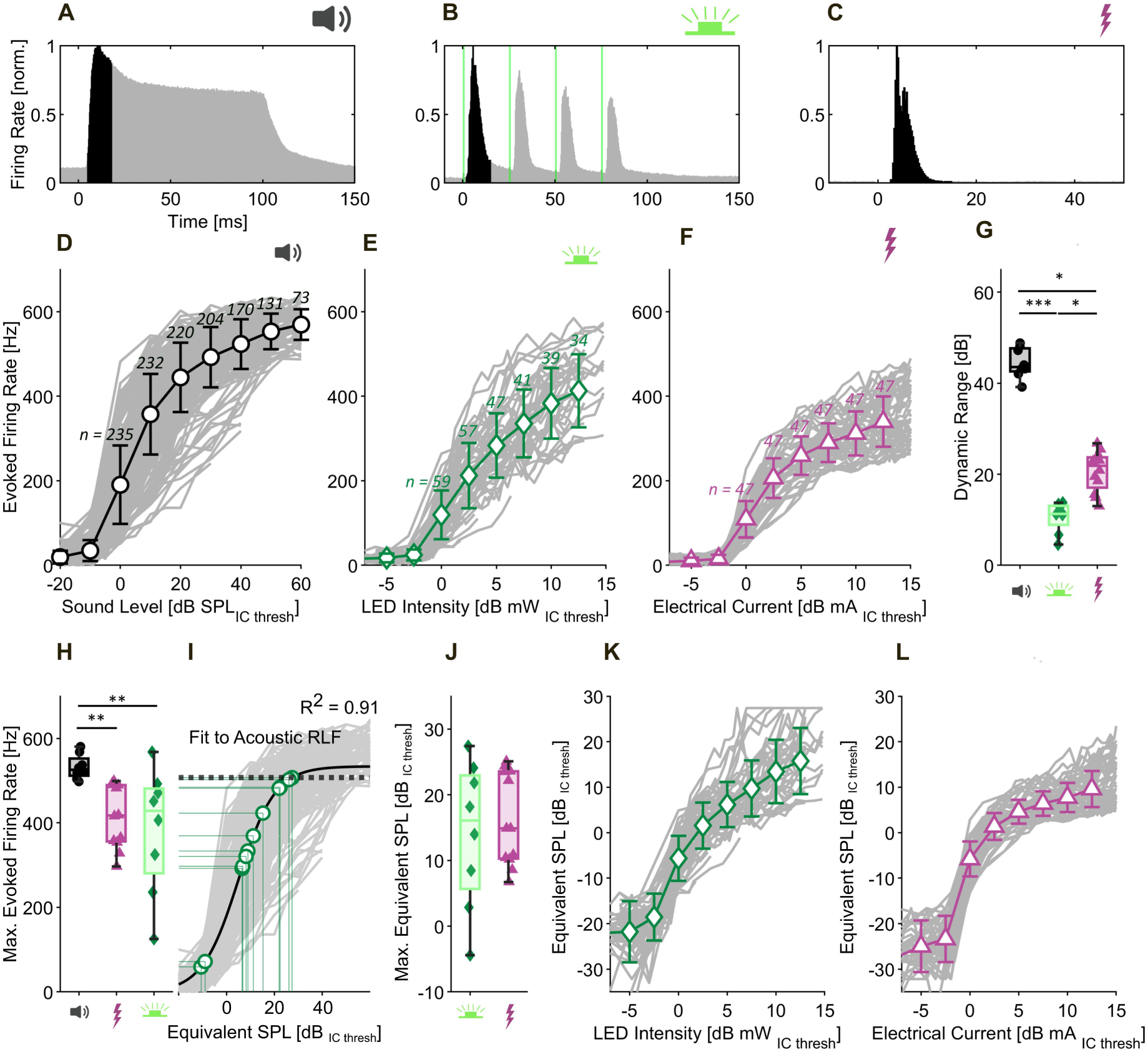
Intensity coding by acoustic, optical and electrical stimulation. **(A-C)** Peristimulus time histograms (PSTHs) in response to a 100 ms tone (**A**), optical pulse train (1 ms, 40 Hz pulse trains, 200 ms inter-stimulus interval, **B**) and electrical pulse (200 µs, ∼3 Hz, **C**). The time windows used to calculate firing rates are highlighted in black. PSTHs were derived by summing and normalizing responses across all recording electrodes to maximum stimulus intensities acquired for each stimulus modality. **(D-F)** Evoked firing rate quantified for each pure tone frequency or optical/electrical emitter using the maximally responding electrode at each stimulus intensity relative to threshold (mean ± SD) for acoustic (**D**), optical (**E**), and electrical (**F**) stimulation. Traces for individual recordings are plotted in the background. Because multiple frequencies or emitters were tested within individual animals, the number above each error bar indicates the number of recordings contributing to that data point. The same maximum absolute stimulus intensity was used across recordings within each modality; therefore, recordings with higher thresholds covered a smaller suprathreshold intensity range and did not necessarily reach saturation. Maximum stimulus intensities were 80–90 dB SPL for acoustic, ∼3 mW for optical, and ∼0.5 mA for electrical stimulation. **(G)** Distribution of dynamic ranges for acoustic, optical and electrical stimulation across animals. Recordings obtained from the same animal were averaged before plotting. **(H)** Distribution of the maximum evoked spike rate achieved by acoustic, optical, and electrical stimulation. Recordings obtained from the same animal were averaged before plotting. **(I)** Quantification of an equivalent sound pressure level (SPL) relative to the pure tone ICC threshold for optical and electrical stimulation. A sigmoid fit to the rate-level function for acoustic stimulation (**D**) (pseudo-R² =0.91) was used as a reference to map firing rates elicited by optical and electrical stimulation onto an equivalent SPL. The mapping procedure is illustrated by thin green lines projecting the maximum optical firing rate onto the acoustic rate-level function and from there to the corresponding stimulus intensity in dB SPL. Firing rates exceeding 95% of the fitted asymptote (black dashed line) were censored at the corresponding 95% value before SPL conversion to avoid inflated equivalent SPL estimates caused by extrapolation near the saturating part of the fit. **(J)** Comparison of the maximum equivalent SPL, derived from the mapping in **H**, achieved by optical and electrical stimulation. Recordings obtained from the same animal were averaged before plotting. **(K-L)** Mapping of evoked firing rates achieved by optical (**E**) and electrical stimulation (**E**) to an equivalent SPL (mean ± SD). The data shown in **E-F** was mapped onto the rate-level function for acoustic stimulation as shown in **I**. Traces for individual recordings are shown in the background. The number of recordings contributing to each data point corresponds to the values shown in **D-F**. The same recordings were used for analyses in panels **D–L** (acoustic: n = 236 recordings from N = 8 gerbils; optical: n = 59 recordings from N = 8 gerbils; electrical: n = 47 recordings from N = 12 gerbils). Across the figure, acoustic data are shown in black, optical in green, and electrical in purple. Data for electrical stimulation was reanalyzed from (Dieter et al., 2019). Statistical significance is indicated by stars (*p < 0.05, **p < 0.01, ***p < 0.001). Groups were compared using a Kruskal–Wallis’s test followed by Tukey’s post hoc comparisons after averaging datapoints obtained from the same animal. Exact p-values are reported in **Table EV1**.cBoxes in **G**, **H**, **I** and **L** indicate quartiles and median, with whiskers extending to the minimum and maximum values. Throughout the figure, acoustic data are shown in black with round markers, optical data in green with diamond markers, and electrical data in purple with triangle markers. Icons in **A-G**, **H**, **J** were created in BioRender. Albrecht, N. (2026) https://BioRender.com/c5wtvp1 (speaker), https://BioRender.com/f3676t6 (LED), https://BioRender.com/uo7uoed (electric).

Across stimulation modalities, rate-level functions showed that the ER increased with stimulus intensity and tended to saturate at higher intensities (**Fig. 2D–F**). To compare rate-level functions across recordings with different thresholds, stimulus intensity was related to threshold intensity and plotted in decibel using a factor of 20 for acoustic: dB (SPL) and electrical stimulation: dB (mA), whereas a factor of 10 was used for radiant flux: dB (mW). Dynamic range was quantified as the stimulus range spanning 10-90% of the maximal ER across all 32 recording electrodes (**Fig. 2G**). Saturation was reliably reached in acoustic and electrical stimulation, but not consistently for optical stimulation within the tested intensity range. Because the same maximum light intensity (∼3 mW) was applied across experiments, recordings with higher optical thresholds covered a smaller range of suprathreshold intensities, resulting in fewer recordings at the highest relative intensities (**Fig. 2E**). Importantly, recalculating the rate-level functions using only recordings with data available at every plotted suprathreshold intensity yielded similar rate-level function shapes across in modalities (**Appendix Fig. S3**).

Given the lack of obvious firing rate saturation for optical stimulation, we also calculated the stimulus range covering 10-100% of the maximum ER for optical stimulation, which increased the dynamic range estimate from 11.3 dB for the 10-90% of max ER estimate to 14.2. dB (**Appendix Fig. S4**).

The maximum ER differed between stimulus modalities and was larger for acoustic stimulation than for optical or electrical stimulation (p = 0.004 and p = 0.002, respectively; Fig. 2D–F, H). We mapped the maximal ER of optically and electrically evoked ICC activity onto the rate-level function of pure tone stimulation and related the result to the average pure-tone threshold measured in our dataset (35 dB, Appendix Fig. S5, Fig. 2I). This comparison indicated that a single LED achieved a median maximal ICC activity level equivalent to ∼51 dB (SPL) of pure tone stimulation (Fig. 2H-J). In comparison, the maximal activity elicited by individual eCI electrodes was equivalent to ∼50 dB (SPL). By mapping the firing rates achieved at each step of the optical and electrical rate-level function onto the acoustic reference, each optical and electrical stimulus intensity relative to ICC threshold was assigned an equivalent SPL relative to the pure tone threshold (Fig. 2K-L).

### Spectrally confined tonotopic activation of the spiral ganglion by multichannel oCI

We then characterized the spectral spread of cochlear activation for individual LEDs of multichannel oCI (5 or 10 LEDs) in comparison to pure tones and previously recorded monopolar eCI stimulation (Dieter et al., 2019). Spatial tuning curves (STCs) were constructed by sorting responses across recording sites and stimulus intensities. STCs (**Fig. 3A-E**) showed tonotopically ordered activation by 10 LED-oCI (**Fig. 3B, C**) and to a lesser extent in the framework of 5 LED-oCI **(Fig. 3D**). We found a tendency of STCs to broaden for LEDs at the high-frequency cochlear base (e.g., see star in **Fig. 3C**). Moreover, in some cases, we found additional peak(s) positioned toward both the higher, but more commonly the lower frequency range relative to the main peak (see arrowhead in **Fig. 3B, D**). Next, we used postmortem phase-contrast X-ray tomography of the implanted cochlea to register LEDs relative to the tonotopic axis of the spiral ganglion, on which we mapped the neural activity predicted from the STCs recorded in the ICC for a given LED (**Fig. 4**). We fixed the oCI using dental cement at the round window niche to avoid oCI extraction. We used a modified Greenwood function to describe the frequency map of the spiral ganglion inside Rosenthal’s canal (colored ribbon next to ganglion in **Fig. 4B**, Methods, Khurana et al., 2022). This analysis indicated that secondary peak of ICC activity likely resulted from the spread of light to the tonotopic range of the ganglion opposite to the primary site of LED stimulation (e.g. arrowhead in 1^st^ panel of **Fig. 4B**). This finding suggests that, at least for a rodent cochlea, off-tonotopic target activation can occur already for radiant flux in the sub-mW range.

**Figure 3.**
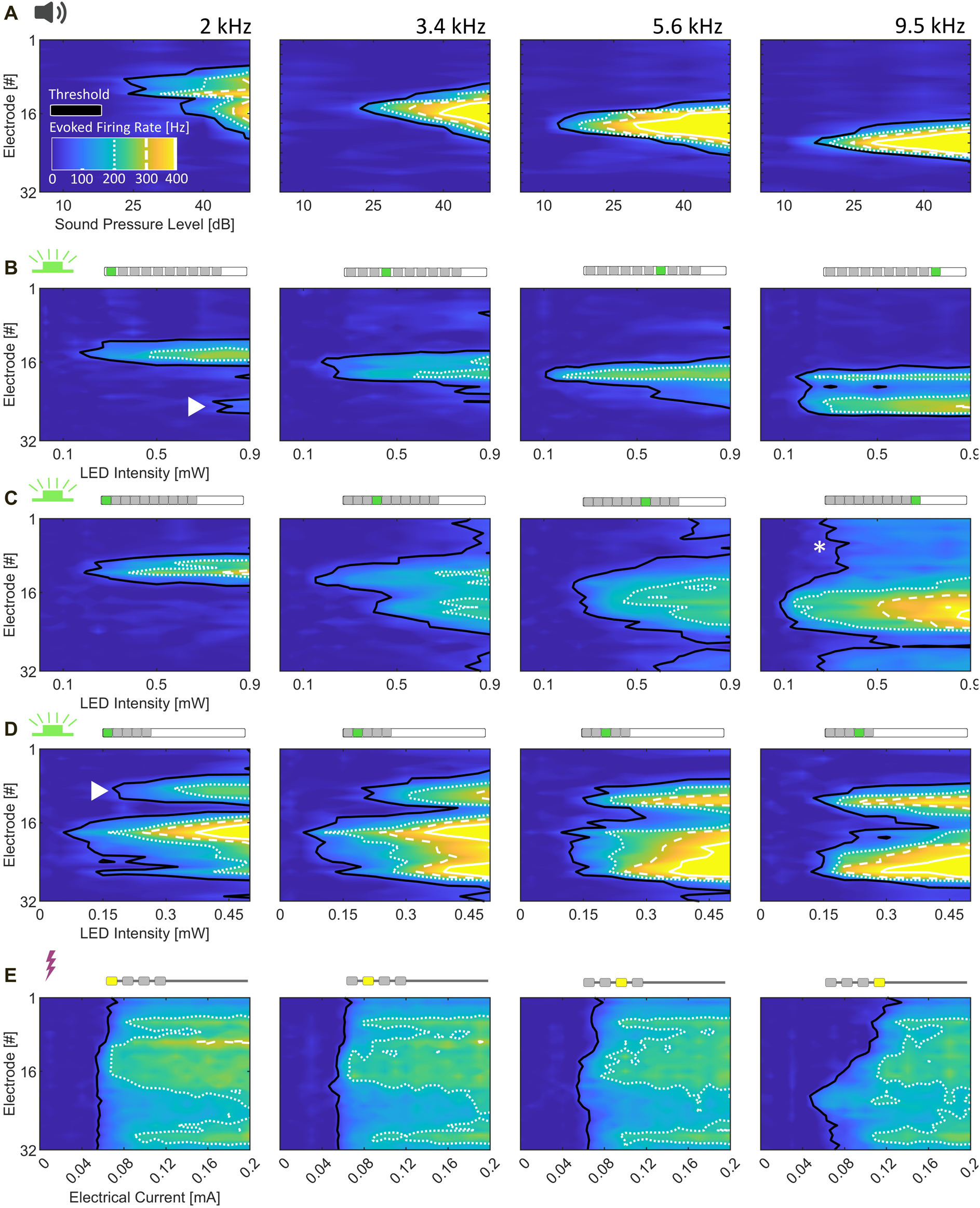
Exemplary spatial tuning curves for cochlear stimulation by different modalities. **(A)** STCs in response to pure tone stimulation with different frequencies in an example animal. **(B-D)** STCs in response to stimulation with individual LED emitters using 1 ms pulses. Each row corresponds to one animal. Columns in **B** show STCs for LED emitters positioned approximately 3600, 2400, 1200, and 0 µm from the round window (emitters 1, 4, 7, and 10, respectively). Columns in **C** show STCs for emitters positioned approximately 2700, 1800, 900, and 0 µm from the round window (emitters 1, 4, 7, and 10, respectively). Columns in **D** show STCs for emitters positioned approximately 1200, 900, 600, and 300 µm from the basal-turn cochleostomy (emitters 1–4, respectively). **(E)** STCs in response to monopolar electrical stimulation with individual electrodes in an example animal. The eCI electrodes were placed approximately 1800, 1200, 600, and 0 µm from the round window (electrodes 1–4, respectively). Data in **E** was reanalyzed from Dieter et al. 2019. Electrode numbers increase from dorsal to ventral recording positions in all STCs shown in **A-E**. The colorbar in **A** is valid for all STCs shown in this figure. The threshold (d’ ≥ 1) is represented by a black contour line, while white contour lines represent evoked firing rates of 200 (dotted), 300 (broken) and 400 Hz (solid). Icons as in **Fig. 2** and pictograms were created in BioRender Albrecht, N. (2026) https://BioRender.com/a2p6ph4 (10-LED oCI), https://BioRender.com/a389tc1 (5-LED oCI), https://BioRender.com/2fyg1kv (eCI).

**Figure 4.**
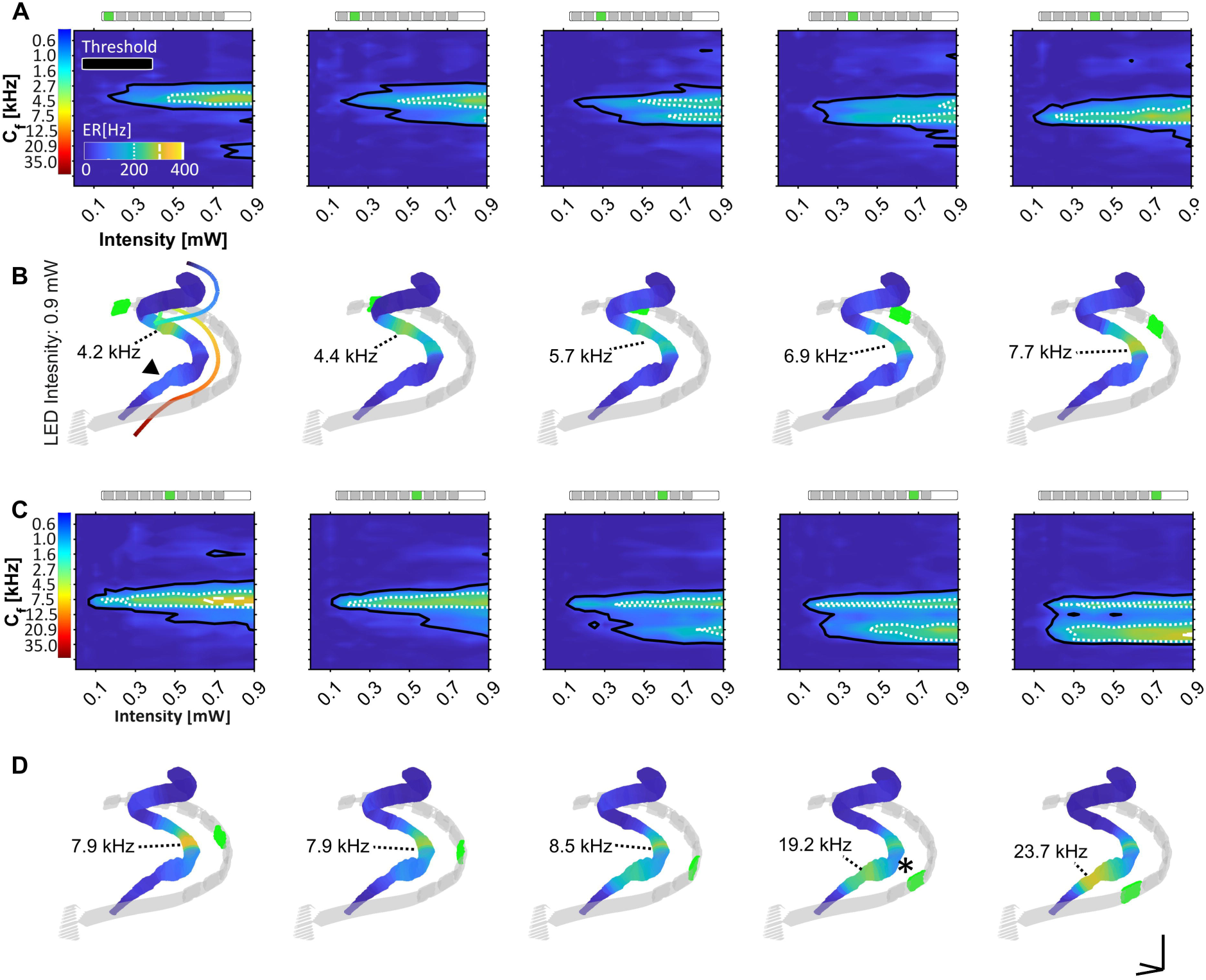
Tonotopic organization of oCI-mediated cochlear activation predicted by ICC activity. **(A, C)** STCs for stimulation with 1 ms pulses from individual LEDs placed along the tonotopically organized Rosenthal’s canal (RC). The color bar indicates characteristic frequency at each recording site derived from pure-tone stimulation prior to oCI implantation. Frequencies outside the gerbil hearing range were not extrapolated. Consequently, not all recording sites were assigned a C_f_. The oCI was implanted through the round window and oriented towards the RC as depicted in **Fig. 1B**. The threshold (d’ ≥ 1) is represented by a black contour line, while white contour lines represent evoked firing rates of 200 (dotted), 300 (broken) and 400 Hz (solid). Pictograms were created in BioRender (see **Fig. 3**). **(B, D)** Projection of the ICC evoked firing rates elicited by 0.9 mW LED radiant flux as shown above (**A**, **C**) onto the RC. The RC and oCI position were reconstructed from X-ray tomography scans of the explanted cochlea (**Fig. 1B**). The active LED is highlighted in green. The strongest responding tonotopic place at 0.9 mW is annotated in each panel (black dashed line). The colored line shown next to the RC reconstruction in **B** corresponds to the colorbar in **A** and represents the expected C_f_ at each RC position according to the gerbil Greenwood function (Müller, 1996). Individual LEDs measured 220 × 270 µm (width × length along the implant). Scale bar in **D** (5^th^ panel) corresponds to 500 µm in each direction.

Moreover, the lateral position and more tangential orientation of the LEDs relative to Rosenthal’s canal within the wider scala tympani at the cochlear base may account for the broader activation observed in several recordings (e.g. 3 to 5 panels of **Fig. 4C, D**; **Fig. 3C**, emitter 10). We initially hypothesized that the pronounced central activity minimum observed in some responses (e.g., star in the 4^th^ panel of **Fig. 4D**) resulted from partial obstruction of the light path by the basilar membrane. However, µCT reconstruction localized penetration of the basilar membrane near LED 7, which did not show such an activity minimum (**Fig. EV3**), arguing against this explanation. Thus, the origin of the minimum remains uncertain with potential reasons including a locally increased impedance of the ICC recording electrode.

Next, we quantified the phenomenon of multiple spectral peaks of activation as well as the tonotopy of optogenetic activation (**Fig. 5**). The spectral peak occurring at the lowest threshold was termed the main spectral peak, while any peaks occurring at higher intensities were termed secondary spectral peaks (**Fig. 5A**). When determining the number of spectral peaks across all recordings of ICC activity elicited by single LEDs (**Fig. 5B**), we found ∼50% of the recordings to contain only the main spectral peak, while one secondary peak, i.e. two peaks in total, were found in ∼50% and two secondary peaks, i.e. three peaks in total, in only two recordings. We estimated the intensity range encoded by the main spectral peak alone (‘Range before Secondary Peaks’, **Fig. 5A**) as the difference between radiant flux at d’ = 1 (threshold) of the lowest threshold secondary peak and at d’ = 1 of the main peak to be 0.60 [0.14, 0.89] mW (median [IQR: 25, 75 percentile], **Fig. 5D**). Moreover, the maximal ER represented in the main peak before the occurrence of any secondary peaks amounted to 272 [235, 306] Hz (**Fig. 5E**). The spectral distance, defined as the difference in characteristic frequency (C_f_) derived from the tonotopic map between the best electrodes of the main and secondary peaks, was 3.1 octaves for secondary peaks at lower frequencies (negative values in **Fig. 5F**) and 4.6 octaves for those at higher frequencies (positive values in **Fig. 5F**).

**Figure 5.**
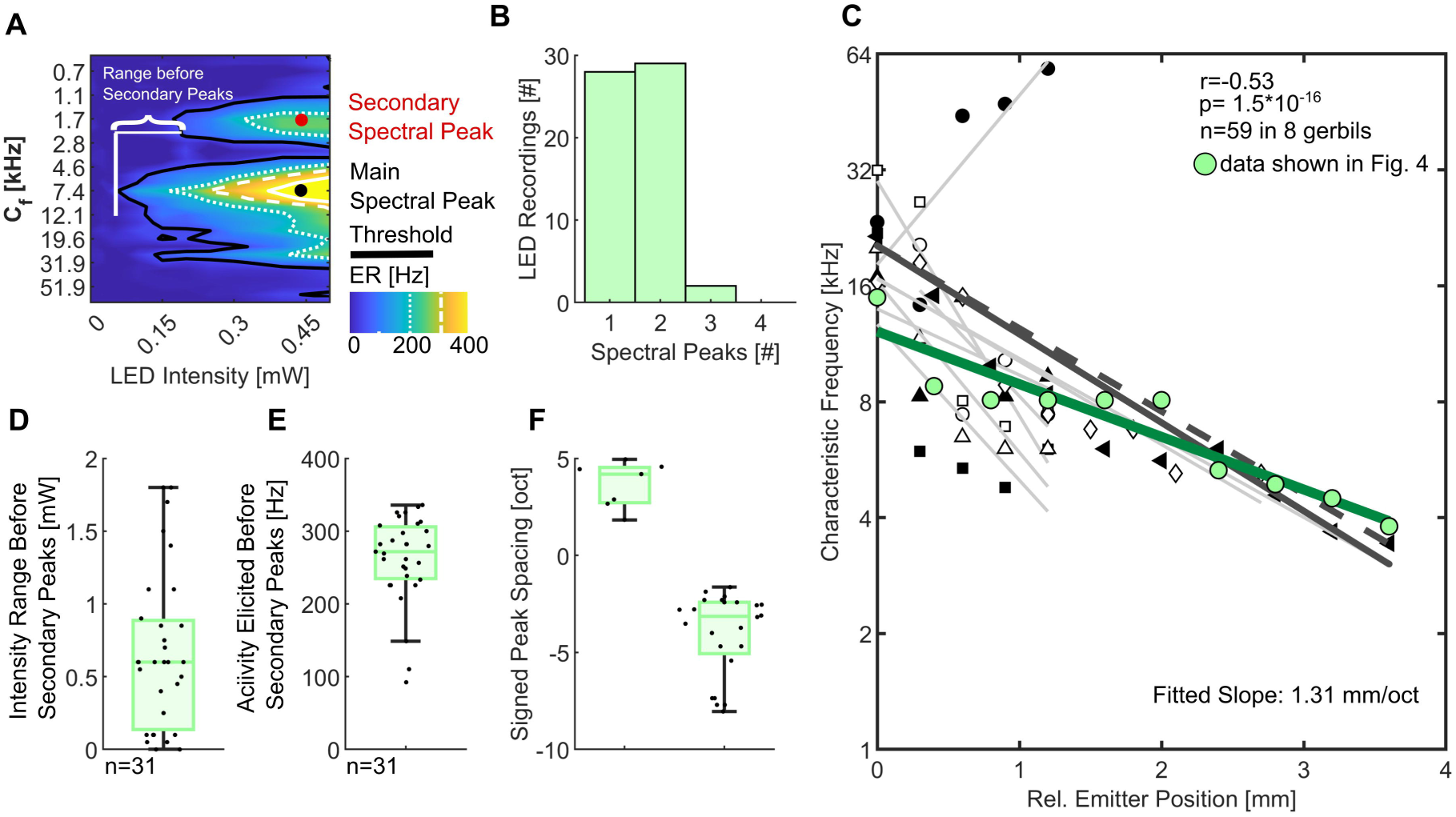
Tonotopic organization of oCI-mediated ICC responses. **(A)** Quantification of response location: Spectral Peaks were automatically detected and assigned to a characteristic frequency (C_f_) based on the C_f_ of the recording electrode with the lowest activation threshold (best electrode). The peak occurring at the lowest threshold was defined as the main spectral peak, while all peaks occurring at higher stimulation intensities were defined as secondary spectral peaks. The threshold (d’ ≥ 1) is represented by a black contour line, while white contour lines represent evoked firing rates of 200 (dotted), 300 (broken) and 400 Hz (solid). **(B)** Histogram of ICC recordings showing one, two, three or four spectral peaks in response to stimulation with individual LEDs. **(C)** C_f_ at the best electrode of the main spectral peak plotted as a function of emitter position relative to the most basal channel. Each combination of marker type and color corresponds to ICC recordings from an individual animal. Grey lines show regression fits for individual animals, while the solid dark line shows the fit across animals r=0.53, p=1.5*10, n=59 LEDs in 8 gerbils). The green colored line indicates the linear fit to the recordings shown in Fig. 4, with the corresponding raw data overlaid as filled green-colored circles. The dashed line indicates the expected slope based on the gerbil Greenwood function for frequencies >2 kHz (1.4 mm/oct; Müller, 1996); its intercept was adjusted to the average fit to facilitate visual comparison of slope differences. **(D)** Intensity range encoded by a single spectral peak in response to simulation by individual LED emitters. The range was calculated as the difference between the thresholds of the main spectral peak and the next occurring spectral peak. **(E)** Activity elicited in a single peak without the occurrence of any secondary spectral peaks, quantified as evoked firing rate. **(F)** Distance between the C_f_ at the best electrode of the main spectral peak and any secondary peaks. Negative values indicate a higher C_f_ for the main peak, while positive values indicate a lower C_f_ compared to the main peak. Boxes in **D-F** indicate quartiles and median, with whiskers extending to the minimum and maximum values.

We quantified the tonotopic organization of oCI stimulation by correlating the intracochlear emitter position and the C_f_ at the best, i.e. lowest threshold, electrode of the ICC response (**Fig. 5C**). Across animals, the fit between best electrode and emitter location yielded a slope of 1.31 mm/oct, which was close to the slope expected by the gerbil Greenwood function (1.4 mm/oct; Müller, 1996). In most individual animals, the C_f_ at the best ICC electrode shifted from lower to higher frequencies as intracochlear emitter position changed from apex to base. In one outlier animal (dark filled circles in **Fig. 5C**), the relationship was reversed, which may reflect atypical emitter placement, as exact emitter positioning is difficult to verify during the experiment. We did not find evidence for systematic differences in ICC threshold in response to stimulation by individual LEDs along the tonotopic axis (**Appendix Fig. S6**).

Next, we investigated the spectral spread of cochlear activation upon stimulation by single LEDs as a measure of spectral selectivity in comparison to acoustic and electrical stimulation (Dieter et al., 2019; Middlebrooks and Snyder, 2007). To do so, we quantified the spatial extent of activation along the tonotopic axis in the ICC at fixed activity levels defined by the maximum ER evoked by a given stimulus (100-350 Hz; **Fig. 6A-C**). Specifically, the width of the STC was defined as the distance between the dorsal-most and ventral-most electrodes within the ICC threshold contour (dT = 1), measured at the lowest intensity level that elicited each ICC activity level. This conservative estimate included all activity regardless of the number of spectral peaks but see below for analysis of the main spectral peak. We subsequently converted this electrode-based measure into spectral spread (in octaves) by multiplying it with the median tonotopic slope across our recordings (**Appendix Fig. S1**).

**Figure 6.**
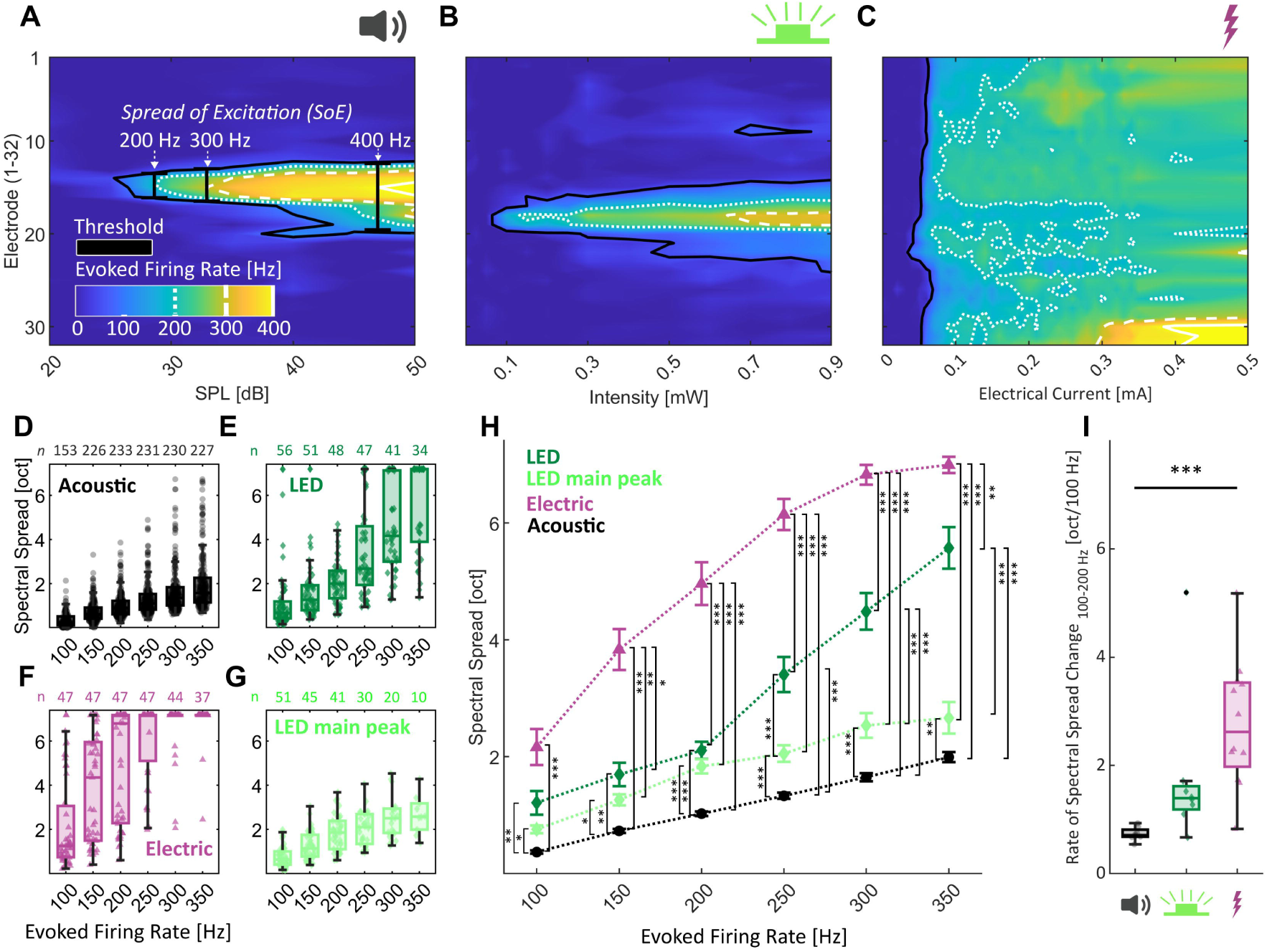
Spectral spread of cochlear excitation for optical, acoustic, and electrical stimulation. **(A-C**) STCs for sound, light and electrical stimulation. Quantification of spectral spread in different stimulation modalities: Spread of excitation (SoE) is measured as the width of the STC at different activity levels quantified as the highest evoked firing rate in response to a given stimulus (example shown for 200, 300 and 400 Hz in **A**). The threshold (d’ ≥ 1) is represented by a black contour line, while white contour lines represent evoked firing rates of 200 (dotted), 300 (broken) and 400 Hz (solid). Icons were created in BioRender (see **Fig. 2**). **(D-G)** Spectral Spread at different evoked firing rates for acoustic (**D**), optical (**E**) and electrical **(F)** stimulation. For optical stimulation the spectral spread was additionally quantified while excluding any secondary spectral peaks (**Fig. 4E**). Because multiple frequencies or emitters were tested within individual animals, the numbers above each panel indicate the number of contributing recordings. **(H)** Comparison of spectral spread (Mean ± SEM) for different stimulus modalities, sound, light and electrical stimulation, as shown in **D-G**. Statistical comparisons were conducted using a linear mixed-effects model, with animal treated as a random factor and evoked firing rate and modality treated as categorical fixed effects, (*p<0.05, **p<0.01, **p<0.001). Evoked firing rate was modeled as a categorical factor to avoid assuming a linear relationship between firing rate and the outcome. Only significant effects are indicated. Model specifics are given in the methods and **Table EV2**. The number of recordings contributing to each activity level is shown in **D–G** and was obtained from N = 8/8/12 gerbils for acoustic/optical/electrical stimulation, respectively. **(I)** Rate of spectral spread change per 100 Hz increase in ER for acoustic, electrical, and optical stimulation. For each animal, the parameter was obtained from a linear fit to the spectral spread values measured at 100, 150, and 200 Hz. Statistical significance was assessed after averaging values obtained from measurements in the same animal using a Kruskal-Wallis test followed by Tukey’s post hoc pairwise comparisons (*p < 0.05, **p < 0.01, *p < 0.001). Boxes in **D-G** indicate quartiles and median, with whiskers extending to the minimum and maximum values. Exact p-values are reported in **Table EV1**. Throughout the figure, acoustic data are shown in black with round markers, optical data in green with diamond markers, electrical data in purple with triangle markers, and the optical main-peak condition in light green with diamond markers.

Previous studies used each animal’s individual tonotopic slope to convert spectral spread to octaves (Dieter et al., 2019; Keppeler et al., 2020). This was intended to correct for lower spread of excitation estimates in animals with higher tonotopic slopes, because a higher slope could indicate a compressed tonotopic representation due to non-ideal oblique placement of the multielectrode array. The correction therefore assumes that tonotopic slope and spectral spread are negatively related. In our dataset they were not: the two were uncorrelated across modalities (Pearson’s r = 0.16, p = 0.41). Thus, we did not apply the correction using each animal’s individual tonotopic slope.

We found that spectral spread increased with activity level, quantified as evoked firing rate (ER), for all stimulation modalities (**Fig. 6D–H**). Across the tested activity levels, the estimated spectral spread was largest for electrical stimulation, smallest for acoustic stimulation, and intermediate for optical stimulation. At low-to-moderate activity levels of up to 200 Hz, the average difference between optical and electrical stimulation increased, reaching a maximum of 2.85 octaves, or 57% relative to electrical stimulation, at 200 Hz. Over the same range, optical stimulation produced significantly greater spectral spread than acoustic stimulation, with a comparatively stable average optical-acoustic difference of 0.8-1.1 octaves, or 70 to 52% of the optical spectral spread. Above 200 Hz, the spectral spread of optical stimulation increased more strongly, narrowing the optical-electrical difference to 1.42 octaves, or 20% of the electrical spectral spread at 350 Hz. Conversely, the optical-acoustic difference increased to 3.58 octaves at 350 Hz. The difference between optical and acoustic as well as electrical and acoustic stimulation was statistically significant at all tested activity levels (p < 0.05, linear mixed-effects model (LME) with Holm correction) while the difference between optical and electrical stimulation was significant for activity levels ≥150 Hz. Thus, optical stimulation provided greater spatial specificity than electrical stimulation while yielding spectral-spread estimates intermediate between electrical and acoustic stimulation.

To determine whether secondary, off-target spectral peaks contributed to the increased spectral spread at higher activity levels, we repeated the analysis after excluding these peaks and quantifying the spectral spread of the main spectral peak of activation. From 250 Hz onward, the spectral spread estimated from all peaks exceeded that of the main peak alone by 1.35–2.9 octaves (40–52% of the all-peak spread; p < 0.05, LME with Holm correction). At 350 Hz, where the opto-acoustic difference was largest, the optical main peak exceeded acoustic stimulation, by 0.68 octaves (25% of the main-peak spread), compared with a difference of 3.58 octaves (64% of the all-peak spread) when all optical peaks were included. Thus, secondary tonotopic activation contributed measurably to the increased spectral spread of optical stimulation, predominantly at higher activity levels.

Because fewer recordings reached the highest activity levels, we repeated the spectral-spread analysis using only recordings contributing across the full tested activity range (**Appendix Fig. S7**). The main conclusions were unchanged, except that the difference between acoustic spread and the LED main peak was no longer significant for activity levels below 300 Hz (p > 0.05, LME with Holm correction). This likely reflects both the reduced sample size (n = 10 LED main-peak analysis) and emphasis of the balanced subset for recordings with lower thresholds and clearly separable main peaks. To facilitate comparison with previous publications (Dieter et al., 2019; Middlebrooks and Snyder, 2007), we additionally quantified spectral spread as a function of dT, a signal detection theory index to compare neural activity levels. This yielded the same ordering of modalities, whereby the optical main peak did not differ significantly from acoustic stimulation (**Appendix Fig. S8**).

To inform future coding strategies, we next quantified the rate at which spectral spread increased with activity level for the range where the benefit of optical stimulation was most visible (up to 200 Hz ER, **Fig. 6I**). For each stimulation modality, we fitted the relationship between spectral spread and evoked response rate up to 200 Hz ER and expressed the slope as the increase in spread per 100 Hz ER. LED-based stimulation showed an increase of 1.38 [1.17, 1.60] octaves per 100 Hz, indicating a slower loss of spectral selectivity with increasing activity level than for electrical stimulation, where spread increased by 2.60 [1.96, 3.50] octaves per 100 Hz. However, at the animal level, this difference was not significant most likely due to the large variance in the estimate for electrical stimulation (p>0.05, **Fig. 6I**). Acoustic stimulation showed the smallest rate of spectral spread change, with an increase of 0.60 octaves per 100 Hz ER, which was significantly lower than for electrical stimulation (p < 0.001, **Fig. 6I**) but not significantly lower than for optical stimulation.

### Addressing channel discriminability for multichannel optical cochlear implants

Discriminability depends, at least in part, on how distinctly different stimulation channels are represented in neural population activity. This was first quantified using representational similarity analysis (RSA), which captures the overall organization of relationships among ICC response patterns within each stimulation modality. For each pair of tones or o/eCI channels, representational dissimilarity (RD) was calculated as the correlation distance between their ICC response patterns at matched activity levels (**Fig. 7B**, Kriegeskorte, 2008; Sabesan et al., 2023). For example, **Fig. 7B** illustrates how the ICC response patterns evoked by two stimuli, i.e. two tones, electrical or optical stimulation channels, are compared to obtain their pairwise RD. First, for each stimulus the lowest intensity eliciting a given firing rate (e.g. 200 Hz in **Fig. 7B**) is identified. RD is then calculated as the correlation distance between the responses across all 32 ICC electrodes at these matched activity levels.

**Figure 7.**
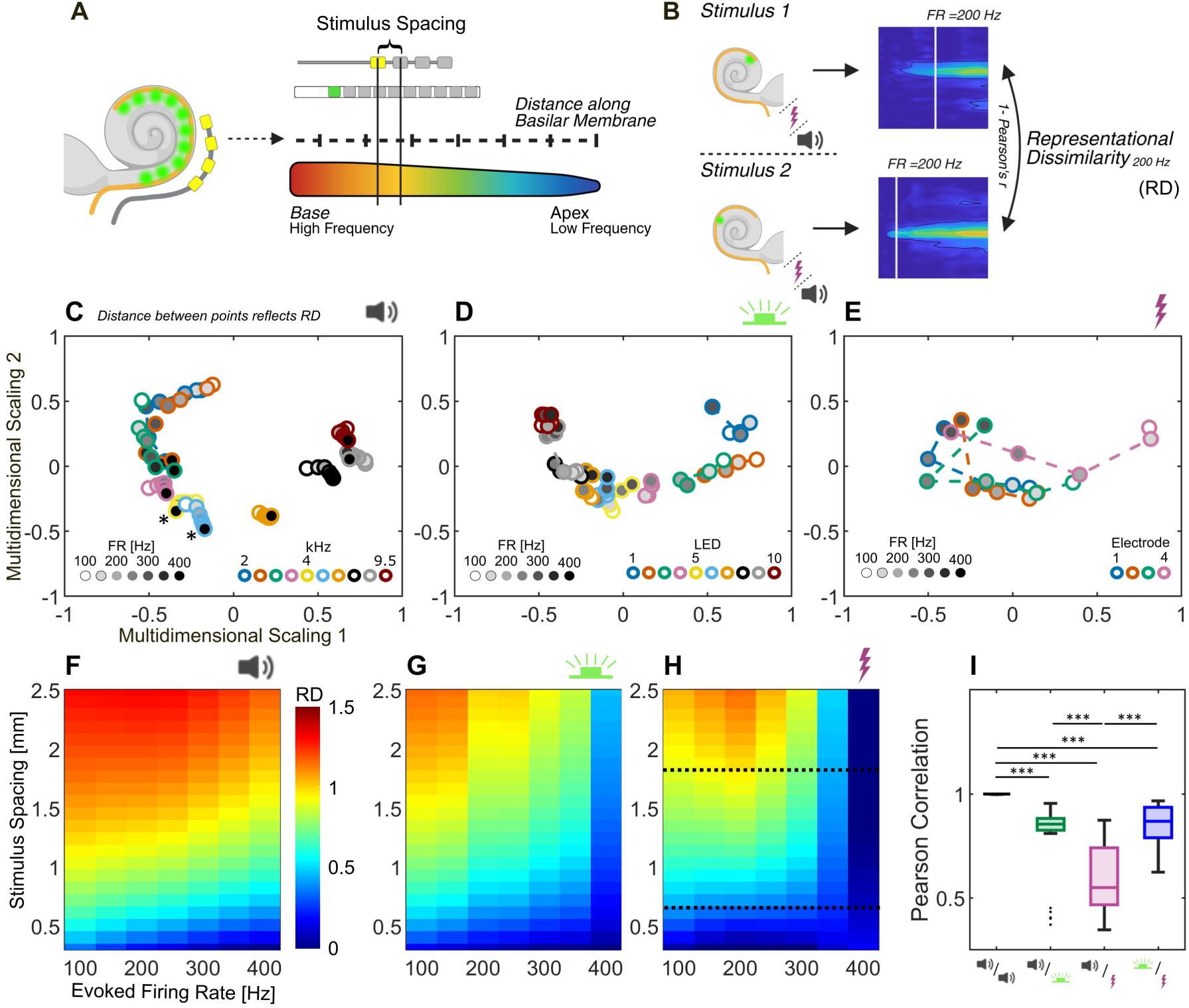
Representational similarity analysis indicates more physiological response patterns for multichannel optical cochlear implants. **(A)** Greenwood-mapped stimulus spacing across modalities. Physical spacing between LED or electrode pairs was compared to the expected cochlear activation spacing elicited by pairs of pure tones, calculated using a factor of 1.4 mm/oct which corresponds to the gerbil cochlear place-frequency map (Müller, 1996). Created in BioRender. Albrecht, N. (2026) https://BioRender.com/835lm6f. **(B)** Quantification of representational dissimilarity. Responses to pairs of stimuli (two tones, optical pulses, or electrical pulses) were compared by computing the correlation distance (1 – Pearson’s r) across the 32 recording electrodes. Responses were matched by activity level, defined as the maximum ER elicited by each stimulus (example shown in **B**). Representational dissimilarities for all pairs of LEDs, electrodes, or tones within a single animal were visualized using multidimensional scaling (**C-E**). Created in BioRender. Albrecht, N. (2026) https://BioRender.com/plry1a8. **(C-E)** Visualization of example representational geometries for acoustic (**C**), optical (**D**), and electric (**E**) stimulation in individual animals. Multidimensional scaling was used to represent the correlation distance between pairs of stimuli, such that the distance between data points reflects their correlation distance. Color coding indicates different frequencies (**C**), LED emitters (**D**), or electrodes (**E**), while the marker filling denotes activity level measured as achieved firing rate. **(F-H)** Visualization of average representational geometries for each stimulation modality (columns). Each matrix cell shows response dissimilarity for a given stimulus pair as a function of activity level (abscissa). Average dynamics were predicted using a linear mixed-effects model across stimulation modality, stimulus spacing, activity level, and their interaction as well as animal as a random effect. Extrapolated regions were present for the eCI data and lie outside the black dashed lines (**H**). **(I)** Distribution of similarity in signal dynamics across stimulation modalities. The model used for **K–M** was applied to predict signal dynamics for each individual animal. Pearson’s correlation coefficients were computed between matrices for all pairs of animals to assess similarity across modalities. As described above, this analysis was restricted to the activation spacings for which data were available in all modalities (600–1800 µm). Statistical differences were evaluated using a Kruskal–Wallis test followed by Tukey’s post hoc comparisons. Acoustic data are shown in black, optical in green, and electrical in purple. Data for electrical stimulation were reanalyzed from Dieter et al., 2019. Statistical significance is indicated by stars (*p < 0.05, **p < 0.01, ***p < 0.001). Exact p-values are reported in **Table EV1**. The number of recordings contributing to each activity level is shown in **Fig. EV4** and was obtained from N = 8/8/12 gerbils for acoustic/optical/electrical stimulation, respectively. Icons in C-I were created in BioRender (see **Fig. 2**).

Because acoustic frequency differences are naturally represented along the cochlear tonotopic axis, the evoked representational dissimilarity (RD) patterns provide a biologically relevant reference against which artificial stimulation can be compared. RSA was therefore applied to test whether the within-modality pattern of pairwise RD, the representational geometry, among optical or electrical stimulation channels resembles the corresponding pattern among acoustic frequencies. To enable this comparison, frequency spacing, typically measured in octaves, was matched to the physical spacing of e/oCI channels using the Greenwood function (**Fig. 7A**).

To visualize the organization of pairwise RD, we projected the responses of representative animals into two-dimensional space using multidimensional scaling (MDS; **Fig. 7C–E**). In these plots, responses with lower RD are positioned closer together, whereas larger distances indicate more dissimilar ICC response patterns. For acoustic stimulation, MDS revealed distinct clustering of responses evoked by different tones that were largely preserved across higher stimulation intensities (**Fig. 7C**). For example, responses to 3.3, 4, 4.8, and 5.7 kHz, indicated by pink, yellow, blue, and orange circles, respectively, remained largely separated across activity levels. Activity level is indicated by the circle filling. The separation between 3.3 and 4 kHz became particularly apparent at higher activity levels (stars in **Fig. 7C**). Similar trends were observed for optical stimulation, where responses to different LEDs clustered in a similar way (**Fig. 7D**). For eCI, in contrast, responses evoked by the same electrode at different activity levels differed more strongly than responses evoked by different electrodes such that clear channel-specific clusters were not preserved (**Fig. 7E**). This is visible as responses to different electrodes, indicated by differently colored circles, shifting together from right to left with increasing activity level rather than remaining grouped by electrode identity.

To quantify these effects across animals, we modeled RD as a function of stimulus spacing for each modality and activity level using LMEs (**Fig. 7F-H, Fig. EV4, Table EV2**). For acoustic stimulation, RD increased with tone spacing and did not diminish at increasing stimulus intensities. At lower activity levels, these trends were closely resembled by optical stimulation, while at higher stimulation intensity responses to different LEDs became less distinct. Electrical stimulation showed an even stronger reduction in RD with increasing stimulus intensity. Accordingly, at higher activity levels (250, 350, 400 Hz), optical stimulation produced RD that were significantly more comparable to those evoked by pure tones than did electrical stimulation (p < 0.05, **Fig. EV4**). Finally, comparison of the overall representational geometry across stimulus modalities showed that oCI-evoked activity more closely resembled acoustic representational geometry than eCI-evoked activity did (p = 0.0008, **Fig. 7F-I, Table EV2**). Notably, this comparison was restricted to the spacing range sampled in all three modalities (600–1800 µm; black dashed lines in **Fig. 7H**).

While RSA captures the relative structure of pairwise response dissimilarities across modalities, it does not directly measure how accurately individual response patterns can be discriminated by a downstream decoder. We therefore turned to linear discriminant analysis (LDA) to quantify discriminability more directly. Using cross-validated decoding at matched activity levels and corresponding stimulus spacings (**Fig. 7A-B**, **Fig. 8A**), we assessed how well responses evoked by two different tones or stimulation channels could be separated on a trial-by-trial basis. Pairwise decoding accuracy was then related to stimulus spacing to obtain neurometric functions for each modality (**Fig. 8B-E**, **Fig. EV5**). Across modalities, decoding accuracy generally increased with increasing stimulus spacing, although this trend was less evident for electrical stimulation because only a restricted spacing range (600–1800 µm) was available (**Fig. 8C-E**). In contrast, the effect of activity level differed markedly across modalities. For acoustic stimulation, decoding accuracy increased with activity level (**Fig. 8C**). Optical stimulation showed a similar improvement up to approximately 250 Hz, followed by an apparent decline at higher activity levels (**Fig. 8D**). Electrical stimulation, by contrast, showed a progressive decrease in decoding accuracy with increasing activity level (**Fig. 8E**). Notably, the same qualitative differences were also apparent within the spacing range sampled in all three modalities.

**Figure 8.**
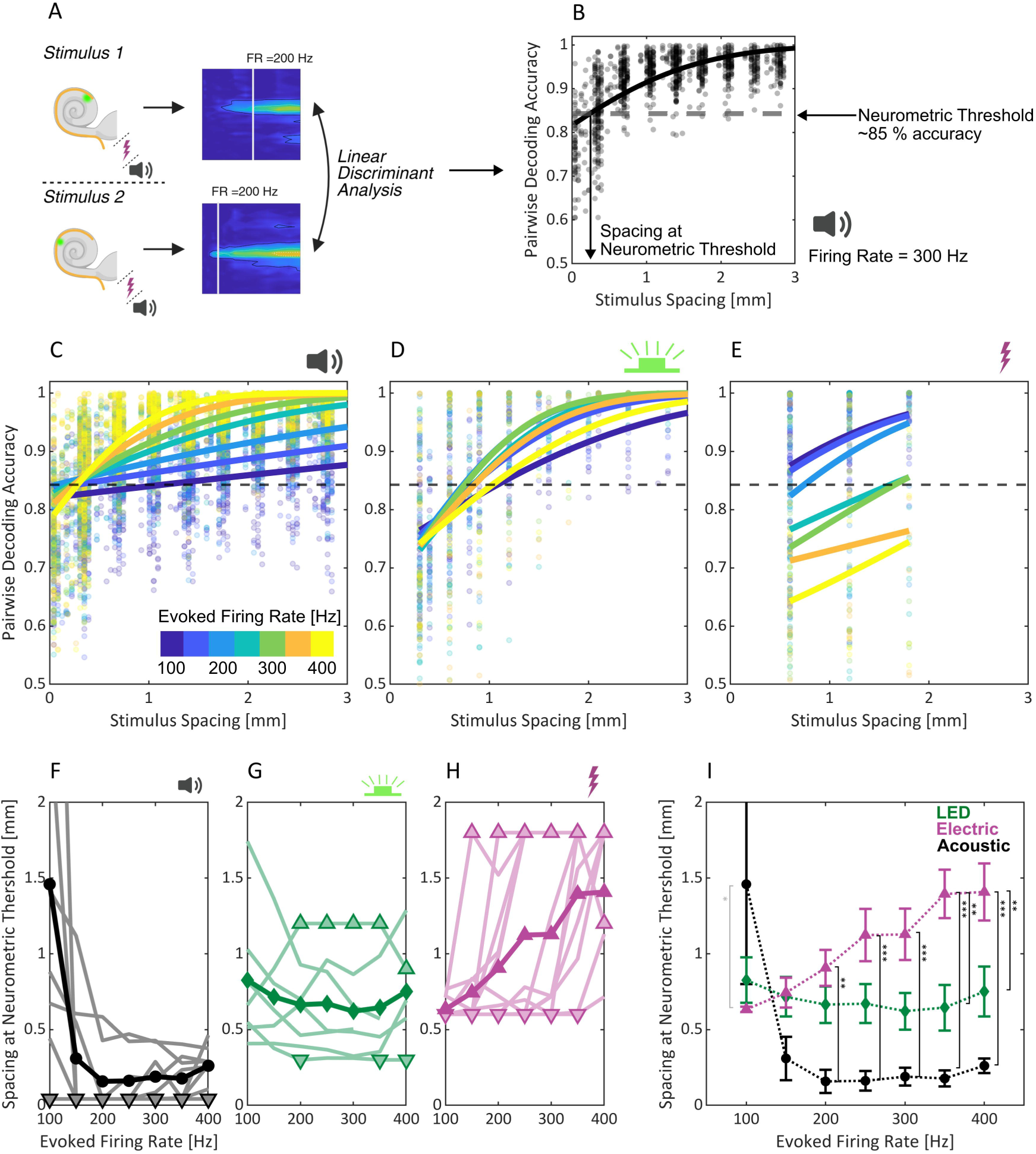
Linear discriminant analysis reveals robust channel discriminability across firing rates in multichannel optical cochlear implants. **(A)** As in Fig. 7, we quantified the discriminability of responses to stimulus pairs (two tones, optical pulses, or electrical pulses) now using linear discriminant analysis. Responses were matched by activity level, defined as the maximum ER elicited by each stimulus (example for 300 Hz in **B**). Classifiers were trained on 30 trials per stimulus, and performance was estimated using 5-fold cross-validation repeated 50 times with different random trial partitions; accuracy was then averaged across repetitions. Created in BioRender. Albrecht, N. (2026) https://BioRender.com/6p8ne8s. **(B)** Neurometric functions (solid black line) based on a cumulative normal two-alternative forced choice model were used to visualize decoding accuracy as a function of stimulus spacing. A common threshold criterion (arrow) was derived from the acoustic condition (84.3 %), and the spacing at which each fitted curve reached this criterion was used as a measure of spectral resolution across modalities. **(C–E)** Neurometric functions across activity levels (ER) for acoustic (**C**), electric (**D**), and optic stimulation (**E**). Different colors denote different evoked firing rates, as indicated in **C**. Background points show the decoding accuracy for individual stimulus pairs at each ER. **(F–H)** Spacing at the neurometric threshold crossing (**Fig. 8B**) as a function of evoked firing rate for acoustic (**F**), optical (**G**), and electrical (**H**) stimulation. Pale background lines show individual animals, and darker solid lines show the across-animal mean. Thresholds falling outside the sampled spacing range were clamped to the highest or lowest tested value and are marked by upward or downward triangles, respectively. The number of recordings contributing to each activity level is shown in **Fig. EV5**. Icons in B-H were created in BioRender (see **Fig. 2**). **(I)** Comparison of spacings at the neurometric threshold crossing (Mean ± SEM) for different stimulus modalities i.e., sound, light and electrical stimulation as shown in **F-H**. Statistical comparisons were conducted using a linear mixed-effects model, with animal treated as a random factor and evoked firing rate and modality treated as categorical fixed effects (for specifics see Methods and **Table EV2**, *p<0.05, **p<0.01, **p<0.001). Statistical results at 100 Hz activity level are indicated in grey as these results could not be confirmed in a non-parametric sensitivity analysis. Evoked firing rate was modeled as a categorical factor to avoid assuming a linear relationship between firing rate and the outcome. Only significant effects are indicated. Exact p-values are reported in **Table EV1**. Data were obtained from n = 8/8/12 gerbils for acoustic/optical/electrical stimulation, respectively. Throughout the figure, acoustic data are shown in black with round markers, optical data in green with diamond markers, and electrical data in purple with triangle markers.

For a direct quantitative comparison, we next extracted neurometric threshold estimates from the fitted neurometric functions for each modality and activity level. The threshold criterion was defined from the acoustic dataset as the decoding-accuracy level at which the fitted acoustic neurometric curves varied least across activity levels, yielding a criterion of 83.9 % decoding accuracy (**Fig. 8B**). Consistent with preceding observations from psychophysical experiments (e.g., Micheyl et al., 2012; Wier et al., 1977), the spacing required to reach this criterion decreased in a saturating manner with increasing activity level for acoustic stimulation: a trend likely driven by decreasing trial-to-trial variability with activity level (**Fig. 8F, I**). The same parameter remained comparatively stable for optical stimulation across activity levels, likely indicating that in this dataset, discrimination of oCI channels was not strongly diminished by increasing spectral spread or secondary spectral peaks at higher activity levels (**Fig. 8G, I**). For electrical stimulation, on the other hand, the stimulus spacing needed to surpass the threshold criterion increased monotonically (**Fig. 8H–I**).

At 100 Hz, estimated thresholds were lower for both electrical and optical than for acoustic stimulation, but only the electrical–acoustic difference reached significance. Because the acoustic thresholds were highly variable at this activity level and violated assumptions of normality, we assessed the robustness of these differences using a Kruskal-Wallis test. The differences at this activity level were not reproduced in this analysis and should therefore be interpreted with caution. From 150 Hz onward, the estimated thresholds were largest for electrical stimulation, intermediate for optical stimulation, and lowest for acoustic stimulation. Averaged across this activity range, optical thresholds were 0.44 mm lower than electrical thresholds (range 0.03–0.75 mm across activity levels) and 0.47 mm higher than acoustic thresholds (range 0.4–0.51 mm across activity levels). In relative terms, the electrical threshold was on average about 1.7-fold the optical threshold, increasing from near-parity at 150 Hz to roughly 1.9-fold at 400 Hz, whereas the optical threshold was on average about 3.4-fold the acoustic threshold for higher activity levels (≥ 150 Hz), reaching approximately four-fold at 200 Hz (0.66 vs 0.16 mm). While the difference between acoustic and electrical stimulation was statistically significant for activity levels ≥ 200 Hz (p < 0.01), the difference between acoustic and optical stimulation did not reach statistical significance for activity levels ≥ 150 Hz (p > 0.05). Pure tones were sometimes discriminable at the smallest tested spacing (**Table EV3**), limiting the estimation of their true thresholds, and the 32-channel recording array may not fully capture the spatial representation of acoustic responses in the ICC. Therefore, the difference of e/oCI neurometric thresholds to acoustic thresholds might be underestimated.

Most importantly, particularly at higher activity levels (≥ 350 Hz), optical stimulation supported discrimination at smaller inter-channel spacing than electrical stimulation (p < 0.01). Because the electrical condition sampled only a restricted spacing range, electrical neurometric thresholds at higher activity levels were frequently censored at the upper bound of the available range (**Table EV3**). Consequently, the observed differences in this regime are likely conservative and may underestimate the true separation between optical and electrical stimulation. These findings support greater spectral resolution with optical than with state-of-the-art electrical cochlear stimulation.

## Discussion

Optogenetic stimulation of the auditory nerve promises a major improvement of hearing restoration. The increased complexity and risk, compared to the current state of the art eCI rehabilitation necessitates careful benefit-risk analysis. Here, we characterized spectral and intensity coding by multichannel oCI stimulation. We first assessed whether individual optical channels could recruit the auditory pathway over a useful activity range at translationally feasible energy requirements and then scrutinized the discriminability of optical stimulation channels in multichannel oCI using two complementary approaches: representational similarity analysis (RSA, Kriegeskorte, 2008; Sabesan et al., 2023) and linear discriminant analysis (LDA). Enabled by the advent of the powerful ChR ChReef and of optimized LED-based oCI, we find that oCI stimulation of the auditory nerve with a power budget that seems compatible with clinical translation offers sound encoding with spectrally distinct channels.

### Intensity coding by cochlear optogenetics

Clinical translation and preclinical research of optical cochlear stimulation require a close match of channelrhodopsin sensitivity and the oCI’s optical power output (Dieter et al., 2020; Huet et al., 2024; Keppeler et al., 2020). In this context, the threshold and rate-level function results can indicate whether individual oCI channels can evoke sufficiently robust neural responses at clinically feasible light intensities. Responses must be strong enough to support reliable sound encoding, while optical power requirements should remain low enough to limit energy consumption. Achieving this balance is essential, because excessive power demands would compromise battery life, potentially risk phototoxicity and, thereby, challenge safety and practicality of future oCI. By combining ChReef with green LED oCI, we found neural responses in the ICC to stimulation with 0.15 µJ optical pulses (**Fig. 1C**) which compares well with previous work using ChReef (Alekseev et al., 2025). Notably, this energy range is on the same order of magnitude as that used in clinical electrical stimulation (∼0.05 µJ (Zeng, 2017)) suggesting that optogenetic stimulation can approach energy regimes within a range considered feasible for clinical application.

Increasing stimulation energy of individual oCI channels up to 3 µJ evoked increasing firing rates that, in some cases, approached the maximum evoked firing rates achieved with pure tone acoustic stimulation (**Fig. 2D-F**). Yet, median maximal firing rates evoked by single LEDs were lower compared to acoustic stimulation and comparable to maximum firing rates achieved by single eCI channel electrical stimulation (**Fig. 2H**). Compared to acoustic and electrical stimulation, single LEDs rarely reached firing rate saturation (at least at 3 µJ) hampering the estimation of maximal output dynamic range and ICC activation. Hence, the apparent dynamic range of 11.3 dB (mW) likely underestimates the achievable dynamic range. This indicates that further lowering the energy threshold as well as using higher light intensities per channel and/or simultaneous multi-channel stimulation present important avenues for future studies. Lower thresholds could be achieved using ChRs with greater single channel conductance (ChReef: 80 fS; Alekseev et al., 2025), some ChR such as hyperpolarizing KCR1 show 700 fS (Govorunova et al., 2022) and/or membrane expression (Gradinaru et al., 2007; Keppeler et al., 2018). Importantly, with the current demonstration of feasibility of preclinical low-power oCI stimulation, future studies can now evaluate multi-channel stimulation schemes. Using simultaneous multi-channel stimulation schemes for preclinical research will allow charting the range of cumulative ICC activity across frequency bands amenable to oCI which could tentatively inform expectations for loudness perception (Auerbach et al., 2019; Vollmer et al., 2007).

The equivalent sound pressure level approximated for the maximal firing rate achieved by a single LED was 51 dB (SPL). Although this represents substantial neural recruitment by a single optical channel, it remains below sound levels typically encountered during conversational speech (∼65 dB SPL). Importantly, such a comparison is necessarily approximate because the energy of natural sounds is distributed across frequency bands, whereas the present estimate reflects stimulation by a single optical channel. Future studies should address this by multichannel stimulation, as well as by extending the range of neural activity for single oCI channels. The latter could be achieved by AAV-Promotor-ChR combinations enabling higher light sensitivity as well as by greater optical output per oCI channel.

The reduced SGN density observed after transduction may also have contributed to the limited ICC activity by reducing auditory nerve input. As previous studies suggest that this loss may be related to pressure during cochlear injection (Bali et al., 2022), alternative approaches for viral delivery may help preserve SGN density and further increase the achievable neural response range (Thirumalai et al., 2025).

The median equivalent SPL estimate of 51 dB for single LED stimulation was lower than a previous estimate in gerbils for single fiber optogenetic stimulation (67 dB (SPL), (Michael et al., 2023)). Comparing both studies identifies methodological reasons for the discrepancy and offers important insights. Notably, the studies differ in i) ChR choice (CatCh vs. ChReef), ii) optical stimulation strategy (emitter, projection, intensity range), iii) reference acoustic stimulus (click vs. pure tone), and iv) analysis (time window, d’ vs. FR). CatCh required higher light intensities (threshold: ∼2 µJ, range probed: 1-40 µJ or mW) which were delivered from the round window along the modiolar axis for recruiting many SGNs across frequencies (Michael et al., 2023). This is very different from the projection toward fewer nearby SGNs in Rosenthal’s canal from an LED placed in scala tympani in the present study. Previously, broadband clicks were used rather than the now employed tone burst and only the d’ analysis was presented, for which we found an equivalent sound pressure level of 59 dB (SPL) (**Appendix Fig. S9**). Despite these uncertainties, we conclude that appropriate matching of SGN light sensitivity achievable by optogene therapy and light delivery by oCI is of paramount importance and that the current combination indicates substantial coverage of ICC activity even with the upper limits used: 3mW (3 µJ) by single LEDs.

### Spectral encoding by tonotopic and spatially precise stimulation

Findings from focused electrical stimulation showed that improvements in spatial selectivity have often been offset by elevated stimulation thresholds and the need for higher stimulation intensities (Bierer, 2007; Bonham and Litvak, 2008). However, higher stimulation intensities are also associated with broader current spread, which may reduce the effective spatial specificity of stimulation. Against this background, we investigated the spectral coding properties of optical cochlear stimulation across a wide range of firing rates.

A prerequisite for physiologically meaningful spectral encoding by individual oCI channels is their ability to activate SGNs in a tonotopically organized manner. By positioning up to 10 LEDs of the oCI within scala tympani along the tonotopic axis, we observed a systematic shift in the tonotopic location of ICC responses corresponding to the tonotopic position of the stimulating LED (**Fig. 3B-D**, **Fig 4**, **Fig. 5C**). This spatial shift (1.31 mm/oct, **Fig. 5C**) was in good agreement with predictions based on the gerbil Greenwood function (∼1.4 mm/oct, Müller 1996). A previous study employing LED stimulation of CatCh-expressing SGNs reported a smaller spatial shift, which only partially met the expectations from the Greenwood function as well (∼2.54 mm/oct for tonotopic slope of 4.5 mm/oct in the ICC, Dieter et al., 2020).

The spread of excitation (SoE) is a well-established parameter for the spectral resolution of different stimulation modalities (Dieter et al., 2019; Middlebrooks and Snyder, 2007; Snyder et al., 2004). We found the SoE for LED-based stimulation to be significantly lower than for electrical stimulation for evoked firing rates of at least 150 Hz while it was significantly larger than the spectral spread evoked by pure tones for the tested activity levels (**Fig. 6H**). This finding is consistent with previous reports suggesting that optical stimulation of spiral ganglion neurons is more spatially confined than electrical stimulation and higher activity levels can be achieved with spatial precision (Azees et al., 2023; Dieter et al., 2020, 2019; Keppeler et al., 2020). Relating these activity levels to equivalent acoustic sound levels provides a more intuitive estimate of the operating range over which optical stimulation improved spectral selectivity. The SoE of optical stimulation was significantly smaller than that of electrical stimulation at 150 Hz, corresponding to an equivalent SPL of 33 dB. The difference was largest at 36 dB SPL equivalent (200 Hz) and remained significant up to 45 dB SPL equivalent (350 Hz). To inform future coding strategies, we additionally quantified how rapidly spectral spread increased with activity level (**Fig. 6I**). This parameter captures the trade-off between increasing neural activation and preserving spectral selectivity, which is central for defining useful operating ranges of artificial stimulation strategies. In the current oCI implementation in the gerbil cochlea, spectral selectivity was best preserved up to approximately 200 Hz, above which off-target tonotopic regions were recruited (secondary spectral peaks). This range therefore appears most relevant for developing coding strategies with the presently available stimulation levels, although higher activity levels will likely be advantageous for clinical use of optogenetic SGN stimulation. Within this range, spread increased more slowly for optical than electrical stimulation, although this difference was not significant at the animal level, and still faster than for acoustic stimulation.

At higher LED intensities, ICC responses occurred at multiple positions along the tonotopic axis, resulting in what we term secondary spectral peaks (**Fig. 3-5**). Secondary spectral peaks were observed in around half of the recordings of LED-evoked ICC activity (**Fig. 5B**). The secondary peak accounted for much of the broadening of SoE at higher optical intensities (i.e., for ER > 150 Hz). On the other hand, when focusing on the main spectral peak of LED-based stimulation (i.e., excluding secondary peaks from the analysis), the spread of excitation remained lower and more closely followed that evoked by pure tones (**Fig. 6H**). This seems relevant, as we consider the secondary peaks to arise from off-target projection beyond the nearby SGNs in the portion of Rosenthal’s canal facing the LED, i.e. the activation of SGNs on the opposite side of the cochlear turn (cross-turn activation, **Fig. 4**).

Notably, secondary spectral peaks have been observed in previous studies employing cochlear optogenetics, although less frequently, as well as in studies using infrared stimulation of the cochlea (Dieter et al., 2019; Richter et al., 2011). The use of a more light-sensitive ChR ChReef may have increased the likelihood that even low levels of scattered light elicited unintended neural activation. Limiting light emission to the range selectively recruiting the nearby SGNs still enabled a substantial dynamic range (**Fig. 5D-E**). In addition, the off-target activation will be less likely in larger cochleae with higher bone density such as in humans and was not found in a previous modeling study of oCI stimulation of the 2.5-fold larger cochleae of humans (Khurana et al., 2022).

Beyond reducing the spread of individual channels, the functional benefit of optical stimulation ultimately depends on whether different channels activate sufficiently distinct tonotopic populations. Because the ICC is a major midbrain hub through which much ascending auditory information is relayed, distinct ICC population responses to different stimulation channels provide a central neural basis for downstream perceptual or behavioral discrimination (Drotos and Roberts, 2024). A key question was therefore whether the relationships among ICC responses evoked by optical or electrical channels mirror the organization observed for acoustic frequencies, known to evoke distinct responses (**Fig. 7**). We therefore compared the within-modality pattern of responses quantified as representational dissimilarity (RD) across acoustic, optical, and electrical stimulation. This revealed a central advantage of oCI: optical channels showed a more tone-like organization of ICC responses, with RD increasing with channel distance and remaining relatively stable across activity levels (**Fig 7C-H, Fig. EV4**). By contrast, eCI responses were more strongly shaped by activity level than by electrode identity, resulting in weaker channel-specific organization (**Fig. 7E, H**). Specifically, we found optically evoked responses to be more similarly organized to sound-evoked responses for activity levels of 250, 350 and 400 Hz compared to electrically evoked responses (**Fig. EV4**).

To assess whether this more tone-like response organization translated into more directly measurable channel discriminability, we next used linear discriminant analysis (LDA, **Fig. 8**). Consistent with the RSA, LDA showed that optically evoked ICC population responses allowed more reliable pairwise discrimination of stimulation channels than electrically evoked responses, particularly at higher activity levels (**Fig. 8**). Thus, the more structured within-modality organization observed for oCI was accompanied by improved decodability of channel identity from ICC activity. Interestingly, this improvement became significant at higher activity levels (≥350 Hz), although the spectral spread of optical responses also increased at these levels. This suggests that oCI responses can remain meaningfully distinct despite broader overall activation. One possible explanation is that a strong, spatially localized response focus remains embedded within a broader activation pattern (e.g. LED 10 in **Fig. 3C**).

This comparison was based on the relative organization of response similarities and dissimilarities within each modality, rather than on absolute similarity between acoustic, optical, and electrical response patterns. While this limits conclusions about how closely oCI reproduce natural acoustic representations, it also reflects a relevant feature of prosthetic hearing: the goal may not be to recreate acoustic activity patterns exactly, but to provide sufficiently structured and separable neural input that downstream auditory circuits can interpret and adapt to through plasticity. Ultimately, behavioral studies will be required to determine whether the improved spectral encoding observed at the physiological level translates into enhanced perceptual discrimination.

Despite this improvement in channel discriminability, optical stimulation still produced a larger spread of excitation, and neurometric thresholds tended to be larger than for acoustic stimulation (**Fig. 6**, **Fig. 8**). One factor likely limiting spatial specificity is emitter size. The LEDs used in this study had a comparatively large surface area of 220 × 270 μm. To close the remaining gap between acoustic and optical stimulation emitters with smaller dimensions, e.g. optical waveguides, may therefore lead to more targeted illumination of SGNs and improved spectral resolution (Khurana et al., 2022). We observed that the shape of RC activation depended on the orientation of LEDs to anatomical structures: LEDs positioned more tangentially towards the RC in the larger scala at the base of the cochlea produced broader activation patterns (**Fig. 4**). Future implant designs should therefore enable precise, reproducible placement and orientation of the emitters, for example, through a design that flexibly conforms to the cochlear geometry, e.g. avoiding scala crossing, yet positions each emitter oriented orthogonally toward the RC.

### Limitations of the present study

In this study, we investigated the spatial representation of different modalities of cochlear stimulation in the ICC to evaluate the utility of oCI for encoding spectral information. To this end, we quantified firing rates evoked by single optical or electrical pulses and compared them with onset responses to pure tones, thereby emphasizing spatial response patterns rather than temporal coding properties. This matches the different modalities at the level of the analysis window. Future studies could instead match them at the level of the stimulus, for example by comparing single light pulses with brief tone bursts (e.g., 10 ms duration with 5-ms onset and offset ramps). As it will likely be advantageous for clinical hearing restoration, future studies should also focus on faithfully reproducing the sustained temporal pattern of acoustic stimulation, which will require opsins with faster kinetics (Koert et al., 2026).

ChReef’s closing kinetics (*τ*_off_ ∼30 ms at physiological temperature) limits optogenetic coding to stimulation rates up to 50 Hz (Alekseev et al., 2025, accompanying paper Koert et al.). Although studies using low-pass-filtered acoustic or vocoded speech suggest that speech perception is possible with limited temporal-envelope information (Xu et al 2005, Xu and Zheng 2007), particularly at high spectral resolution, the kinetics of ChReef challenges its use for future clinical hearing restoration. Ideally, the gain in spectral resolution should not be traded in by poor temporal coding in oCIs. ChRs with faster temporal kinetics would likely support a more complete representation of auditory signals Indeed, employing fast ChRs such as f-Chrimson enables activation of the auditory system with high spectral resolution while additionally supporting temporal fidelity to a few hundred Hertz (> 200 Hz, Koert et al., 2026)). However, higher activation thresholds challenge the power budget available for clinical application. Future developments should therefore focus on the development of ChRs that combine low activation thresholds with sufficiently fast closing kinetics (*τ*_off_ ≤ 15ms).

Another limitation is that we characterized stimulation by individual oCI channels, whereas clinical sound coding relies on multichannel stimulation. Although interactions between simultaneously activated channels are expected to be lower for optical than for monopolar electrical stimulation (Azees et al., 2023), this cannot be inferred from the present experiments. Addressing channel interaction is relevant given the secondary activation peaks observed at higher stimulation levels in the present study of the small rodent cochlea, which could enhance channel interaction in oCI if off-target stimulation also occurred in the larger human cochlea. Future studies should therefore directly investigate simultaneous multichannel stimulation (Middlebrooks and Snyder, 2007).

Optical and electrical stimulation data in the present study were obtained in the presence of intact inner hair cells. While future clinical application would primarily be aimed at patients lacking functional inner hair cells, previous studies of cochlear optogenetics suggest that deafening does not substantially alter the measures most relevant here (Alekseev et al., 2025; Dieter et al., 2020; Thirumalai et al., 2025). Specifically, activation thresholds and spread of excitation have not been reported to differ significantly between hearing and deafened animals.

Finally, the electrical stimulation data were acquired in a previous study using the same experimental setup, protocol, and analysis pipeline. Yet, the comparison of these datasets remains a limitation, as subtle experimenter-related differences cannot be fully excluded.

## Methods

### Animals

Data were obtained from 27 (15 AAV-injected, 12 non-injected) Mongolian gerbils (*Meriones unguiculatus*) of either sex.

Surgical procedures and electrophysiological recordings were performed under isoflurane anesthesia (5% for induction, 1–2.5 % for maintenance at 0.4 l/min) monitored by breathing rate and absence of hind limb withdrawal reflex. Body temperature was maintained at 37 °C using a heating pad. Analgesia consisted of subcutaneous meloxicam (5 mg/kg) in adult animals or carprofen (5 mg/kg) for postnatal surgeries, and buprenorphine (0.1 mg/kg) administered 30 min prior to surgery and every 5-6 h thereafter.

Animals had ad libitum access to food and water and were housed under a 12 h light/12 h dark cycle. All animal procedures were approved by the local animal welfare authority (Lower Saxony, Germany) and conducted in accordance with national and institutional guidelines.

### Adeno-associated virus

To achieve effective and comparable transduction of SGNs with ChReef we used the same AAV construct in all injected animals. The final vector encoded the ChRmine variant (T218L/S220A) ChReef fused to EYFP and trafficking/export signals under control of the human synapsin promoter and was packaged into AAV PHP.S for intracochlear injection (PHP.S_hSyn-ChRmine(T218L, S220A)-TS-EYFP-ES (ChReef) _WPRE_SV40pA at 2.40E+13 GC/ml (CdPCR)) (Alekseev et al., 2025).

The AAV-construct was produced and purified as previously (Alekseev et al., 2025; Huet and Rankovic, 2021). Briefly, HEK-293T cells were triple-transfected with helper plasmids (TaKaRa), cis-plasmids encoding for ChReef under control of the human synapsin promoter, and trans-plasmids encoding the AAV capsid. Viral particles were harvested from cell lysates and culture supernatants, purified by iodixanol gradient ultracentrifugation, concentrated, and formulated in phosphate-buffered saline. Viral genome titers were determined by quantitative PCR or digital PCR using a commercial AAV titration kit. Vector purity was routinely assessed by silver staining following gel electrophoresis. Purified viral stocks were stored at −80 °C until use.

### Postnatal intracochlear injection

For the injected cohort, 1 µl of the virus construct were injected into the coclear scala on postnatal day 8 or 9 via a transbullar approach after carefully spreading the surrounding tissue (Huet et al., 2021a; Huet and Rankovic, 2021). The cartilaginous bulla was identified using the adjacent facial nerve as a landmark. A glass pipette was then advanced toward the cochlea until perilymph reflux was observed, confirming intracochlear placement before injection of the AAV construct. To provide sufficient time for transduction, gerbils were kept at least to an age of two months before conducting further experiments.

### Acoustic stimulation

Open-field acoustic stimulation was delivered via a loudspeaker positioned 30 cm in front of the animal (Scanspeak Ultrasound; Avisoft Bioacoustics). The system was calibrated prior to experiments using a ¼-inch microphone (51BF-1) and preamplifier (12 AQ; GRAS). Acoustic stimuli consisted of 100 ms pure tones (5 ms cosine on/off ramps) spanning frequencies from 0.5 to 32 kHz and intensities from 0 to 90 dB SPL. Stimulus generation and presentation were controlled by custom-written MATLAB scripts and delivered via a data acquisition card (NI PCIe-6323; National Instruments). Each stimulus was presented 30 times in randomized order with an inter-stimulus interval of 200 ms. In non-injected gerbils the acoustic input of the right ear was blocked by the aid of a 0.25 % Poloxamer hydrogel injected into the right bulla through the right external tympanic duct. This way, the acoustic input was restricted to the left ear in the control group, matching the side of oCI stimulation in injected animals. Acoustically evoked responses were recorded using ABR as well as ICC recordings (e.g. **Fig. 3A, Appendix Fig. S1**).

### Laser-based optical stimulation

For optical stimulation, access to the left bulla was obtained via a retroauricular incision with careful removal of overlying muscle layers. A bullostomy was performed to expose the round window. Following perforation of the round window membrane, an optical fiber (200 μm core diameter, 0.39 numerical aperture; Thorlabs) coupled to a 522 nm green laser (L1C-522, combined with a motorized Power Attenuator (MPA); Oxxius) was inserted into the round window and oriented toward the cochlear apex. Radiant flux was calibrated using an optical power meter (PM103USB, S140C; Thorlabs), and stimulation intensities were determined by linear interpolation between calibrated values. Laser-evoked responses were recorded using ABR as well as ICC recordings (e.g. **Fig. EV1**, **Fig. 1C**).

### Implantation and stimulation with optical cochlear implants

Multichannel oCI were assembled from 5 or 10 LED chips (C527TR2227-S0600; Cree; 220 × 270 μm) emitting at 527 nm, integrated at a pitch of 300 or 400 μm on polyimide substrates and encapsulated in silicone (Keppeler et al., 2020). The cochlea was accessed via the bullostomy used for laser-based stimulation, and oCI were carefully implanted through a basal turn cochleostomy or the round window (**Fig. 1B**). One animal in which the oCI was implanted through a mid-turn cochleostomy was excluded from further analysis to ensure comparability across experiments.

oCI were operated using custom-built stimulation hardware (StimBox) based on the same hardware design as the previously published MobileProcessor and running the NSV1 embedded software (Jablonski et al., 2025). Pulse trains consisting of 1 ms pulses were delivered at various stimulation rates and intensities. Each stimulus was presented 30 times in randomized order with an inter-stimulus interval of 200 ms. LED-evoked responses were captured in ICC recordings (**Fig. 1A**, **C**; **Fig.2-8**)

All oCI channels were calibrated individually using an optical power meter (PM103USB, S140C; Thorlabs). Final stimulus intensities were derived from a quadratic fit to calibrated values.

### Recording of auditory brainstem responses

Hearing ability and transduction success were assessed by recording acoustically evoked auditory brainstem responses (aABRs) in all gerbils and subsequently optically evoked auditory brainstem responses (oABRs) in AAV-injected gerbils (**Fig. EV1**). Optical stimulation was delivered via a laser-coupled optical fiber placed in the round window, as described above.

ABRs were acquired from recording the potential difference between needle electrodes below the pinna and at the vertex at a sampling rate of 50 kHz. A needle electrode at the back of the animal served as the ground.

### Analysis of Auditory Brainstem Responses

Recorded ABR signals were band-pass filtered between 300 and 3000 Hz. ABR waveforms were obtained by averaging responses to 1000 repetitions of the same stimulus. ABR waves were detected semi-automatically using custom-written MATLAB scripts (MATLAB 2024a). Thresholds were defined as the lowest acoustic or optical stimulus intensity that evoked a visually detectable ABR wave (**Fig. EV1B**).

AAV-injected animals exhibiting oABR thresholds below 10 mW radiant flux were subsequently used for recordings of optically evoked inferior colliculus activity; experiments were terminated otherwise (n = 2).

### Recording from the inferior colliculus

Multi-unit activity from the central nucleus of the inferior colliculus (ICC) was successfully recorded in 9 out of 13 AAV-injected animals with oABR-thresholds below 10 µJ and 8 non-injected gerbils, as described previously (**Fig. 1A**, Dieter et al., 2019; Michael et al., 2023; Roos et al., 2026). As described above, one of the injected animals was excluded from further analysis due to the mid-turn cochleostomy.

After head fixation and stereotactical alignment (Luigs-Neumann SM-8) the visual cortex overlying the contralateral ICC was exposed by performing a craniotomy ∼2 mm lateral and ∼0.5 mm caudal of lambda. A 32-electrode linear silicone probe (Acute Probe A1×32-6mm-50-177-A32,177 µm^2^ electrode surface, 50 µm electrode spacing, Neuronexus) was then advanced into brain (∼3.3mm) at a speed of 1µm/s by a micromanipulator to avoid damage to the tissue (Fiáth et al., 2019).

ICC activity was evoked by acoustic or optical stimulation using either laser-coupled optical fibers or LED-based oCI, as described above. A 200 ms inter-stimulus interval was used for all stimulation modalities. Data obtained during eCI stimulation from 12 non-injected gerbils were reanalyzed from Dieter et al., 2019.

Activity of multi-unit clusters was amplified, filtered (0.1/50 to 9000 Hz) and recorded at a sampling rate of 32 kHz using a Digital Lynx SX recording system and the Cheetah v.6 recording software (Neuralynx).

### Detection of multi-unit activity

Multi-unit responses were analyzed using custom-written MATLAB scripts as described previously (Dieter et al., 2019; Michael et al., 2023). Raw data traces were first sorted by recording electrode and aligned by sample timepoints. Before further processing, a global mean for each timepoint was calculated across all electrodes and subtracted from each electrode’s trace (McInturff et al., 2022). As stimulation artefacts typically occur synchronized across recording sites, this effectively minimizes LED-stimulation artefacts. After applying a band-pass filter (0.6 – 6 kHz, 4^th^ order Butterworth filter), the threshold for detection of firing activity was defined as four times the mean absolute deviation within a pre-stimulus baseline (estimated as mean/0.675 for data 100 to 2 ms before stimulation). Threshold crossings were converted to spike times by detecting the nearest maximum. A 1 ms dead time where no further spikes could be detected was implemented after each spike timing to avoid double detection of the same event. For electrical stimulation data, threshold crossings that were detected -0.5 ms to 2.5 ms around the stimulus onset were not regarded as spikes to prevent detection of stimulation artefacts.

### Quantification of activity level

To determine time windows for firing rate calculations, peristimulus time histograms (PSTHs, **Fig. 2A-C**) were derived from responses to 100-ms pure tones (200 ms inter-stimulus interval), 40-Hz light pulse trains composed of 1-ms pulses (100 ms duration, 200 ms inter-stimulus interval), or 200-µs biphasic electrical pulses (100 µs per phase) consistently presented at a stimulation rate of ∼3 Hz. PSTHs were generated by summing responses and normalizing to the maximum presented stimulus intensity (∼3 mW, 80–90 dB SPL, or 500 µA). Neuronal activity was defined as responses exceeding twice the mean baseline activity and first occurred 5.38 ms for acoustic, 2.88 ms for optical and 2.63 ms for electrical stimulation relative to stimulus onset. PSTHs were used to identify response onset and duration for each stimulus modality. To avoid bias introduced by substantially different response durations across modalities, firing rates and all derived parameters (i.e. d’-values) were compared within fixed onset windows (13 ms duration): responses to a single optical or electrical pulse (2–15 ms for optical responses; 2–15 ms for electrical responses) were compared to acoustically evoked activity within the onset window of the response (5–18 ms). Evoked firing rates were calculated by averaging responses across 30 repetitions of the same stimulus and subtracting the mean baseline firing rate measured immediately before stimulus onset in mirrored time windows (−15 to −2 ms for optical and electrical stimulation; −18 to −5 ms for acoustic stimulation).

To determine the threshold for neural activation, firing rates from individual stimulus repetitions were compared with baseline firing rates from the corresponding pre-stimulus windows (**Fig. 1C**, **Fig. 2D-F**, **Fig. 6D-H**). Empirical receiver operating characteristic (ROC) curves were constructed from these distributions, and the corresponding areas under the curve (AUCs) were used to derive dT values via the inverse cumulative normal distribution function (Macmillan and Creelman, 2004; Middlebrooks and Snyder, 2007). The threshold was defined as the first stimulus intensity at which dT exceeded 1. The best electrode was defined as the recording electrode, or the mean of adjacent electrodes, that reached threshold at the lowest stimulus intensity.

### Mapping of characteristic frequencies

Characteristic frequencies (C_f_) were determined for each recording electrode based on responses to acoustic stimulation (**Appendix Fig. S1**). Twenty-five pure tones spanning a frequency range from 0.5 to 32 kHz were presented at sound pressure levels between 0 and 80 dB SPL. For each stimulus frequency, the best electrode was identified. Electrode-specific C_f_ were derived from a linear fit of frequency as a function of the corresponding best electrode and used to establish the tonotopic organization across the electrode array.

### Analysis of Spectral Peaks

Spatial tuning curves (STCs) were constructed by visualizing activity across recording sites and stimulus intensities by drawing iso-contour lines at the threshold (d’=1) and at defined firing rates (200, 300, 400 Hz) using MATLAB’ built-in contour function (**Fig. 3-8**, **Appendix Fig. S10**).

STCs obtained during LED-based stimulation were analyzed by quantifying the number of spectral peaks across the 32 recording electrodes (**Fig. 5**). To minimize subjective bias, we employed a standardized, fully automated analysis pipeline. Threshold contours (d’=1-contourline) were first smoothed along the electrode and intensity axes using a Gaussian window (sigma = 2) to reduce noise-induced local maxima. This smoothing parameter was selected based on an empirical comparison between raw STCs and detected peaks (**Appendix Fig. S10**).

Peaks were identified as local maxima using the MATLAB **findpeaks** function. Where necessary, individual peaks were analyzed separately by partitioning the d’=1-contour at the midpoint electrode between detected peaks. To define the activated area along the tonotopic axis, we determined the best electrode of each spectral peak.

### Spectral spread of excitation

To compare the spread of excitation across modalities, we measured STC spatial width at matched activity levels (**Fig. 6**, Dieter et al., 2019; Keppeler et al., 2020). For each modality, we first found the lowest stimulus intensity that elicited a given activity level (e.g., 200, 300, 400 Hz). STC width was then measured as the distance between the most dorsal and ventral intersections of the threshold contour (i.e., the dT = 1 contour) at that intensity (**Fig. 6A**). Spread of excitation was then converted from distance to octaves by multiplying by a single global tonotopic slope (4.581 oct/mm), defined as the median slope across all animals (n = 28), so that the same conversion was applied to all stimulation modalities.

As an additional parameter, we estimated the rate of change in spread of excitation per 100 Hz increase in firing rate (**Fig. 6I**). This was calculated as the slope of spectral spread as a function of firing rate. For each animal, a linear fit was applied to the spectral spread values measured at firing rates of 100, 150, and 200 Hz, and the resulting slope was multiplied by 100 to express the change in spectral spread per 100 Hz increase in firing rate.

### Representational Similarity Analysis

To quantify response geometries across stimulus modalities, we performed Representational Similarity Analysis (RSA, Fig. 7, Kriegeskorte, 2008; Sabesan et al., 2023). For each animal, we constructed representational dissimilarity matrices (RDMs) from neural responses to different stimuli (e.g., different sound frequencies, LED channels, or CI electrodes) at matched activity levels (**Fig. 7B**). Each RDM entry, referenced as the representational dissimilarity (RD), was defined as the pairwise correlation distance (1 − Pearson’s r) between response vectors, where each response vector consisted of spike rates across 32 recording electrodes. This yielded an **S** × **S** RDM for each animal and activity level, where **S** represents the number of stimulation channels or frequencies tested in that animal. For each RD the corresponding stimulus spacing and activity level were retained for subsequent analyses.

Activity levels were matched as described for spread of excitation. To enable comparison across modalities, stimulus spacing was expressed on a common spatial scale (**Fig. 7A**). For acoustic stimulation, tones spanned 2–32 kHz, similar to the cochlear range covered by the LEDs in the dataset (**Fig. 5**). The physical distance between predicted acoustic activation loci was estimated using a conversion factor of 1.4 mm/oct. According to the gerbil Greenwood function, this approximation is appropriate for frequencies above 2 kHz, where the frequency-to-place conversion changes only weakly, increasing from approximately 1.4 to 1.5 mm/oct across the remaining frequency range (Müller, 1996). These predicted acoustic distances were then matched to the physical spacing of LED channels or CI electrodes. To account for remaining differences in sampling density across modalities, we employed linear mixed-effects (LME) models, allowing valid statistical comparisons despite unbalanced stimulus sets. Prior to model fitting, stimulus spacing was centered around zero and log-transformed using base 2.

First, separate LMEs were fitted for each activity level, predicting RD from modality, centered log-transformed stimulus spacing, and their interaction, while including random intercepts and slopes for stimulus spacing for each animal (**Table EV2**). Statistical comparisons were then performed on per-animal model predictions by computing pairwise areas between predicted curves for each electrical or optical animal and each acoustical animal (**Fig. EV4**). For this comparison only the spacing range sampled in each modality was used.

In a second step, a combined LME was fitted to model RD as a function of stimulus spacing (centered and log-transformed), activity level, and modality, including all interactions between fixed effects (**Table EV2**). Per-animal predictions were compared by computing Pearson correlations between predicted dissimilarity matrices **(Fig. 8F–I**).

### Linear Discriminant Analysis

To quantify decoder-based discriminability across stimulus modalities, we performed linear discriminant analysis (LDA) on ICC population responses (**Fig. 8**). For each animal, modality, and matched activity levels, responses evoked by two different tones or two different stimulation channels were classified pairwise using 30 trials per stimulus. Each trial was represented as a response vector containing evoked spike rates across the 32 recording electrodes. Classification was performed using a linear discriminant classifier with 5-fold cross-validation, repeated 50 times with different fold partitions. Each pairwise comparison thus yielded a cross-validated decoding accuracy reflecting how well the two corresponding response patterns could be discriminated.

For each animal and matched activity level, pairwise decoding accuracies were obtained for all available tone, LED, or electrode pairs, and the corresponding stimulus spacing was retained for subsequent analyses. Stimulus spacing was matched across modalities as described above. Because activation contours were derived from trial-averaged responses, they were not suitable for the trial-by-trial LDA analysis. We therefore identified the stimulus intensity that evoked a maximum firing rate closest to the desired activity level without interpolating by forming contours. Pairwise decoding accuracy was then analyzed as a function of stimulus spacing to derive neurometric functions for each modality and activity level. For this purpose, decoding accuracy was fit with a cumulative normal neurometric function adapted for a two-alternative forced-choice task, with parameters estimated by least squares. Because each decoding-accuracy estimate was based on the same number of trials for each stimulus pair, this fitting procedure was not biased by unequal sampling across comparisons. Neurometric thresholds were defined using a common decoding criterion, selected from the acoustic dataset as the decoding-accuracy level at which the acoustic neurometric curves fitted across animals showed the smallest variation across activity levels (84.3%, **Fig. 8B**). Threshold estimates were then extracted from neurometric curves fitted separately for each animal, as the stimulus spacing at which the fitted curve exceeded this common criterion. This spacing was used as a measure of spatial resolution (**Fig. 8F-I**). When the estimated threshold fell outside the experimentally sampled range, it was censored to the nearest sampled bound (**Fig. 8F-H**, **Table EV3**).**X-Ray tomography**

After completion of recordings and before euthanizing the animal, a subset of oCI were fixed in the stimulation position using dental cement (Paladur, Kulzer). The implant’s position and its orientation relative to Rosenthal’s canal were assessed in explanted cochleae by X-ray tomography, as described previously (**Fig. 1B**, **Fig. 4**, Bartels et al., 2013; Keppeler et al., 2021; Schaeper et al., 2025, 2023; Wrobel et al., 2018). Data acquisition was performed using the micro-CT system ‘Easytom’ (RX Solutions, Chavanod, France) providing a cone beam geometry and equipped with a nanofocus source (Hamamatsu L10711-02, W target, LaB6 cathode, operated at 80 kV) and a CCD camera fibre-coupled to a Gadox scintillator (Ximea, Münster, Germany). Field of view (7.0 x 4.3 mm, width x height) and pixel size (3.5 µm) were adjusted to optimally capture the whole cochlea. 1536 projections were acquired per scan with an exposure time of about 2.8 s resulting in an overall scanning time of approximately 5h. Phase and tomographic reconstruction was performed using the Xact software provided by the manufacturer RX Solutions. Relevant structures, including Rosenthal’s canal and LED emitters, were segmented and visualized using Amira (2019.1, Thermo Scientific) by semi-automated tracing of each component. To visualize ICC activity in the corresponding Rosenthal’s canal (RC) region, we first assigned a characteristic frequency (C_f_) to each RC position by applying the gerbil Greenwood function to the distance along the RC (**Fig. 1B**, referred to as ‘modified Greenwood function; Khurana et al., 2022; Müller, 1996). Electrophysiological responses were then projected onto the RC by matching each ICC recording electrode’s C_f_ to the corresponding position along the RC (**Fig. 4**).

### Immunolabeling and imaging of cochlear sections

Another subset of cochleae (7 of 15 animals) was processed for immunolabeling followed by confocal imaging, as described previously (**Fig. EV2**, Thirumalai et al., 2025).

Fixed cochleae were decalcified for at least 7 days in 0.12 M EDTA (pH 8.0) and embedded in 2% agarose for cross-modiolar vibratome sectioning (VT1200S; Leica; cutting speed, 0.02 mm/s; amplitude, 0.6 mm). Sections of 220 µm thickness were cut at a 90° angle relative to the modiolus to visualize large portions of Rosenthal’s canal within each cochlear turn and to assess potential hair cell transduction. One section per cochlear turn was incubated in blocking and permeabilization buffer (16% normal goat serum, 450 mM NaCl, 0.6% Triton X-100, 20 mM phosphate buffer, pH 7.4).

Primary antibodies against GFP (1:500; chicken; Abcam, 13970), parvalbumin (1:500; guinea pig; Synaptic Systems, 195308), and calretinin (1:500; rabbit; Swant, 7697) were incubated for 72 h at 4 °C followed by three rounds of washing (wash buffer; 20 mM phosphate buffer, 0.3% Triton X-100, 0.45 M NaCl, 3x for 10 min). Secondary antibodies, goat anti-chicken Alexa Fluor 488 (1:200; Invitrogen, A11039), goat anti-guinea pig Alexa Fluor 568 (1:200; Invitrogen, A11075), and goat anti-rabbit Alexa Fluor 647 (1:200; Invitrogen, A21244), were applied for 48 h at 4 °C. After washing twice in wash buffer, twice in PBS, and once in phosphate buffer (5mM), each for 10 min, sections were then mounted and cleared using FOCM (30 % urea, 20 % d-sorbitol, 5% glycerol, dissolved in DMSO, Zhu et al., 2019).

Representative images of Rosenthal’s canal were acquired from each section using a confocal microscope (SP8; Leica) equipped with a 40× objective in immersion oil, with a z-step size of 1 µm.

### Quantification of SGNs in immunolabelled cochlear sections

Cross-modiolar images were analyzed using arivis Vision 4D (Zeiss). Segmentation of spiral ganglion neurons (SGNs) was performed automatically in the parvalbumin channel using a custom model implemented in Cellpose 2.0 (Stringer et al., 2021; Stringer and Pachitariu, 2022). The volumetric boundary for SGN density estimation was computed using a custom-written function to encompass all detected cells. To allow for comparison to 2D-density estimates in previous publications, a representative slice in the middle of the Z-stack was chosen and SGN-density was estimated by dividing the number of cells by the area of the volumetric boundary present in that slice (**Fig. EV2C**). Identification of GFP-positive SGNs was based on the distribution of GFP signal intensity across all cells in the analyzed volume, as described previously, using a Python script implemented within arivis Vision 4D (Huet et al., 2021b; Mittring et al., 2023).

Histological data previously obtained from the left cochleae of nine wild-type gerbils were processed identically and served as a control group (**Fig. EV2B-C**).

### Statistical analysis

Investigators were not blinded to experimental groups. However, most outcome analyses were automated and therefore involved minimal investigator-dependent assessment.

For statistical comparisons, normality of residuals was assessed using Q–Q plots following a one-way analysis of variance (ANOVA) implemented as a linear model, and homogeneity of variances was tested using Levene’s test. As these assumptions were not met for the datasets analyzed, group differences were assessed using the Kruskal–Wallis test for three or more groups followed by MATLAB’s default multiple comparisons test (Tukey), the Mann– Whitney U-test for two unpaired groups, or the Wilcoxon signed-rank test for two paired groups.

For specified analyses, linear mixed-effects models were used to account for repeated-measures structure and unbalanced datasets. Models were fitted in MATLAB using the fitlme function with restricted maximum likelihood (REML). Residual normality was assessed by visual inspection of Q–Q plots and, where necessary, the outcome variable was transformed: a cube-root transformation was applied to the spread of excitation (SoE) in **Fig. 6H**. Final model formulas, as well as coefficient estimates or model performance parameters (for analyses not used for direct statistical inference), are provided in the Appendix. Fixed effects were frequently modeled as categorical factors to avoid assuming a linear relationship between firing rate and the outcome. For predictive analyses, model performance was assessed using root mean squared error (RMSE) and R, computed separately across relevant conditions (**Table EV2**).

### Declaration of generative AI use

Claude Code (Anthropic) was used for programming assistance with the MATLAB analysis and figure code; ChatGPT (OpenAI) was used for coding assistance during initial pipeline development. Claude (Anthropic) and ChatGPT (OpenAI) were used for language refinement and to support literature searches. No generative AI was used to generate or alter data or to derive scientific conclusions. All AI-assisted code and all AI-suggested references were verified by the authors, who take full responsibility for the manuscript.

## Supporting information

Expanded Data View Table 1

Expanded Data View Table 2

Expanded Data View Table 4

## Acknowledgment

We thank Gerhard Hoch, Christiane Senger-Freitag, Ina Preuss, Sarah Schlagowsky, Sandra Gerke, Daniela Gerke, Dr. Christian Gossler, and Ludwig Ehrenreich for expert technical support of the study and Patricia Räke-Kügler for excellent administrative support. We thank Dres. Bettina Wolf, Ursula Fünfschilling, Ramona Brecht and Jana Sie as well as Tabea Stapel for support related to preparing, executing and documenting animal experiments. We thank Dres. Marcus Jeschke, Alexander Dieter and Lakshay Khurana for fruitful discussion on analysis and data. We thank Dres. Fadhel El May (advice on oCI implantation and data analysis) and Sabina Nowakowska (images of cross-modiolar sections of non-injected control cochleae).

This work was supported and funded by Deutsche Forschungsgemeinschaft (DFG) through the Cluster of Excellence (EXC2067) Multiscale Bioimaging EXC 2067/1-390729940 (MBExC, L.R., J.H., T.Ma., T.Mo., T.S.) and the Collaborative Research Center 1690 (K.K., T.Ma., T.Mo., P.R.). The work was further supported by the Else Kröner-Fresenius Foundation via the Else Kröner-Fresenius Center for Optogenetic Therapies (EKFZ, N.A., L.R., J.H., L.J., K.K., T.Ma., T.Mo.) and the European Innovation Council grant OptoWavePro (101158920, T.Mo.). N.A. is recipient of a scholarship of the Goe ngen Promo onskolleg für Medizinstudierende, funded by the Jacob-Henle-Programm/Else Kröner-Fresenius Foundation (2021_EKPK.04) and of the German Scholarship Foundation. A.V. was supported by HORIZON TMA MSCA Postdoctoral Fellowship (OPTOCODE, grant 101107675). E.Ko. is recipient of a scholarship of the Evangelisches Studienwerk e. V. Villigst. N.A., E. Ko., A.V. and L.R. are collegians of the Hertha Sponer College of MBExC and N.A., A.V. and L.R. are members of the EKFZ academy.

## Expanded View Figure Legends

**Figure EV1.**
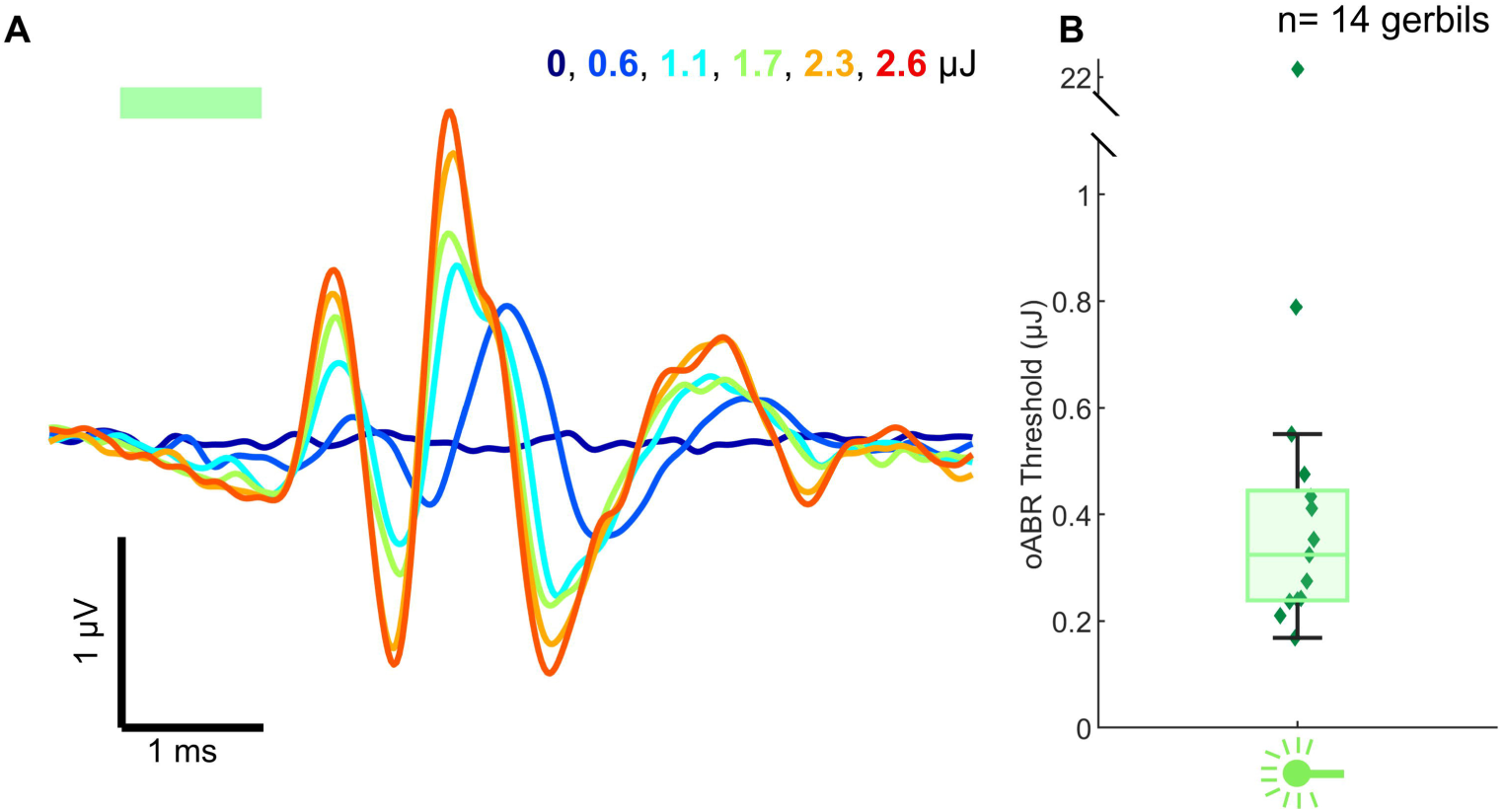
oABRs evoked by laser stimulation. **(A)** Representative oABRs evoked by laser stimulation with an optical fiber positioned at the round window of a Mongolian gerbil cochlea. The fiber emitted 522 nm light, delivered as 1 ms pulses at 17 Hz. Colors indicate radiant energy in µJ. **(B)** Radiant-energy thresholds for the occurrence of oABRs under the stimulus conditions shown in A. n = 14 gerbils. One gerbil did not show oABR responses. Boxes indicate quartiles and medians; whiskers extend to minimum and maximum values. Icon was created in BioRender: Albrecht, N. (2026) https://BioRender.com/mz4pj4h.

**Figure EV2.**
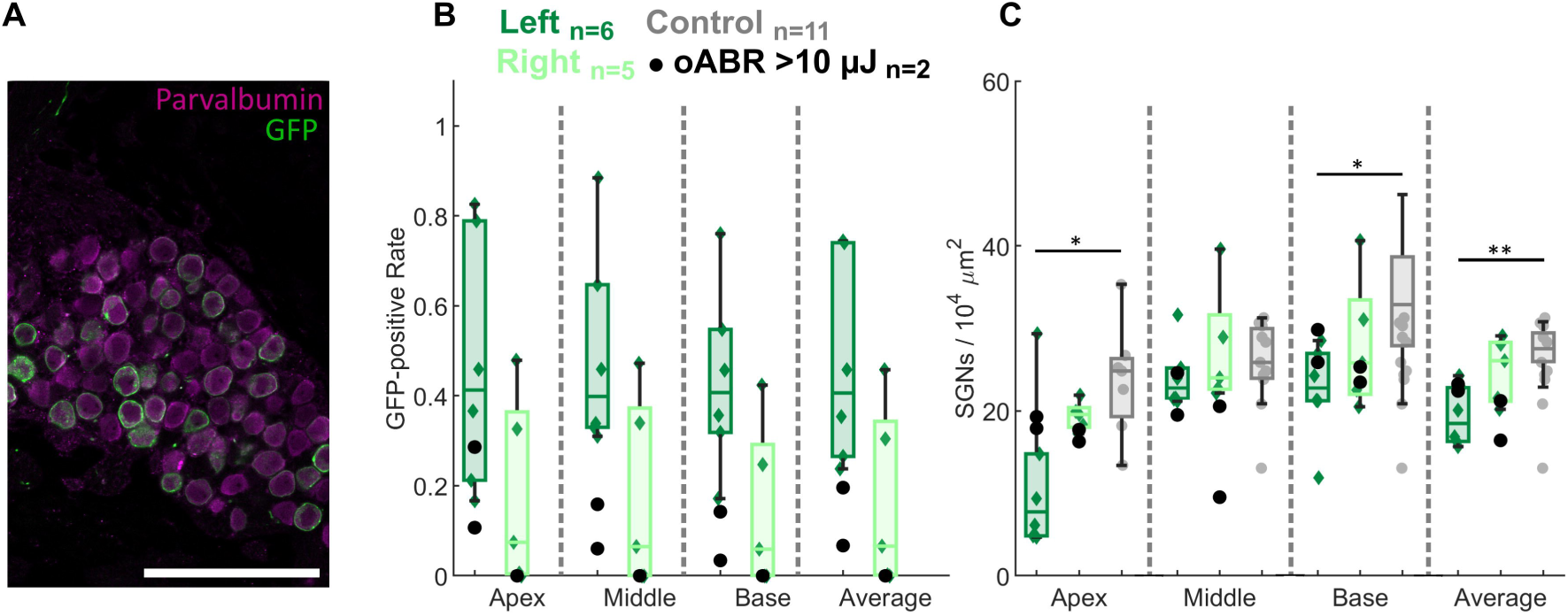
Immunohistochemical Quantification of SGN Transduction Rate and Density. **(A)** Representative confocal image of immunolabeled SGNs in Rosenthal’s canal. SGNs were identified by anti-parvalbumin staining (magenta). Anti-GFP staining was used to label the eYFP tag of ChReef. Scale bar, 100 µm. (**B**) Fraction of GFP-positive SGNs in each cochlear turn as well as averaged across turns of the left, injected cochlea (n = 7) and the corresponding right cochlea (n = 6). No statistically significant difference was detected using a one-sided Wilcoxon matched-pairs signed-rank test (p > 0.05). (**C**) SGN density in each cochlear turn for the groups shown in B, and left, non-injected cochleae from wild-type animals (n = 11). Groups were compared using a Kruskal–Wallis test followed by Dunn’s multiple-comparisons test (* p < 0.05, ** p < 0.01). Only statistically significant differences are indicated. Animals with oABRs detected only at >10 µJ, or with no detectable oABRs, were excluded from statistical testing. Boxes indicate quartiles and medians; whiskers extend to minimum and maximum values.

**Figure EV3.**
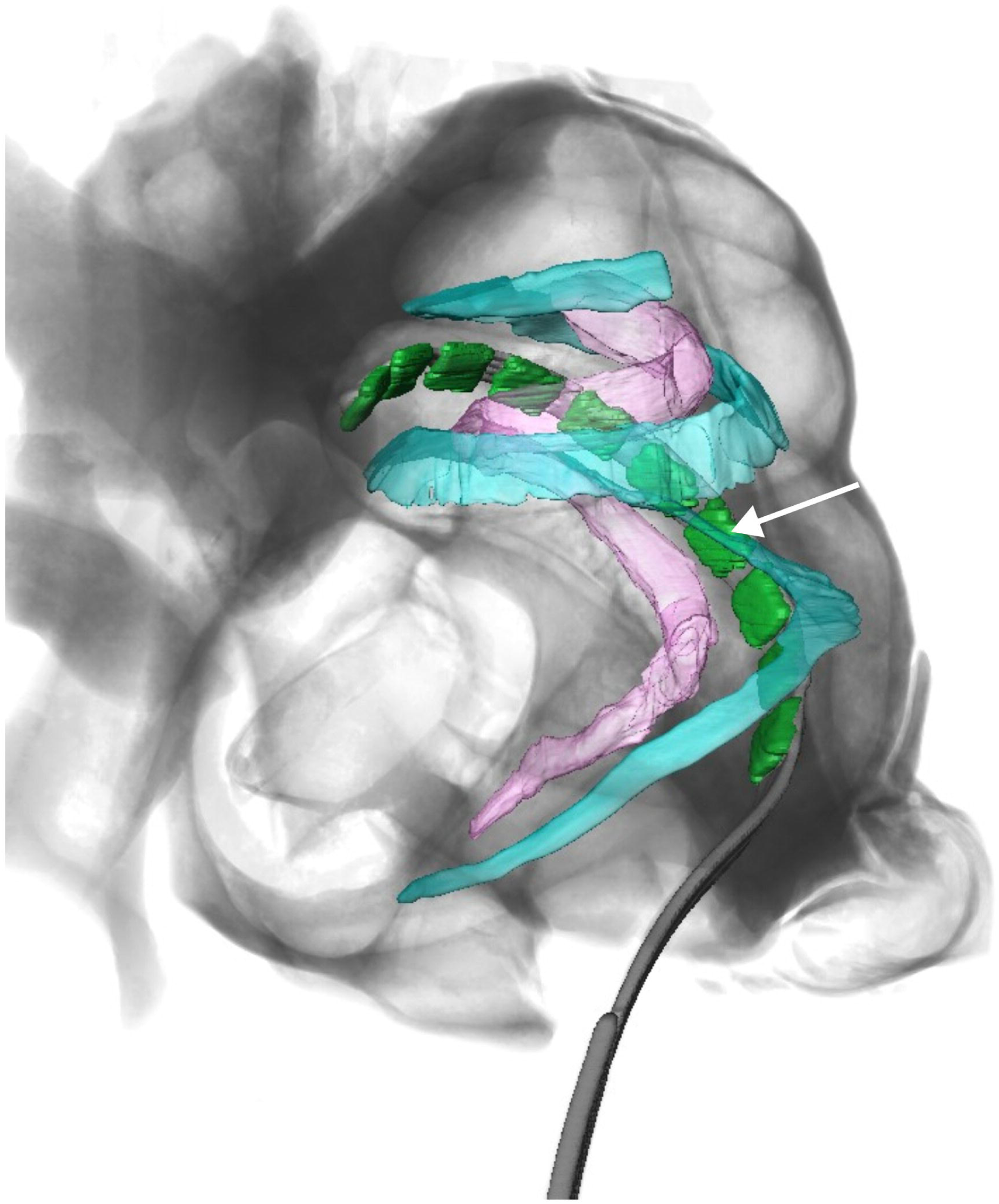
oCI Position relative to the basilar membrane. X-ray tomography of a gerbil cochlea implanted with an oCI, as in Fig. 1B. The oCI pierced the basilar membrane during implantation. In the position used for stimulation, the basilar membrane crossed the implant in front of LED 7 (counted from the apex; white arrow). The estimated positions of the basilar membrane (cyan) and Rosenthal’s canal (magenta) were segmented and visualized after reconstruction. Individual LEDs measured 220 × 270 µm (width × length along the implant).

**Figure EV4.**
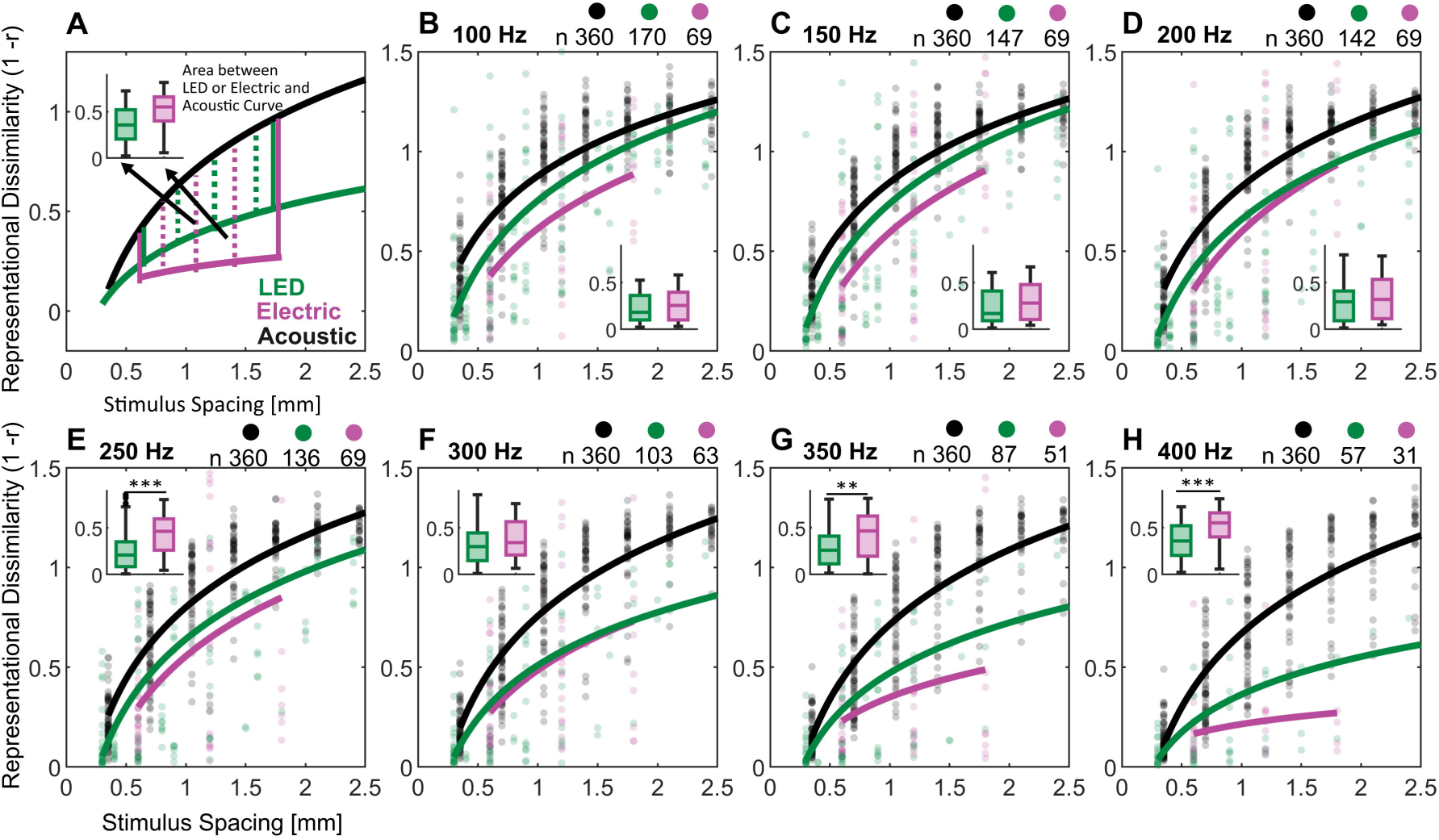
Representational dissimilarity across firing rates. **(A-H)** Comparison of representational dissimilarity across stimulus spacings for different stimulation modalities at matched activity levels (columns). Individual data points, representing representational dissimilarities between stimulus pairs, are shown in the background, and the number contributing to each modality is indicated above the panels. Modality-specific estimates across animals (solid lines) were obtained at each activity level using linear mixed-effects models (see methods, Tab. S2) Differences to the acoustic reference were quantified as the areas between the predicted lines for each animal pair (opto–acoustic and electric–acoustic; method illustrated in Panel F). This analysis was restricted to the activation spacings for which data were available in all modalities (600– 1800 µm). The results are shown as boxplot insets. The significance of differences between oCI and eCI was assessed using a Mann-Whitney U-Test. Only significant differences are indicated (*p<0.05, **p<0.01, **p<0.001). Exact p-values are reported in **Table EV1**.

**Figure EV5.**
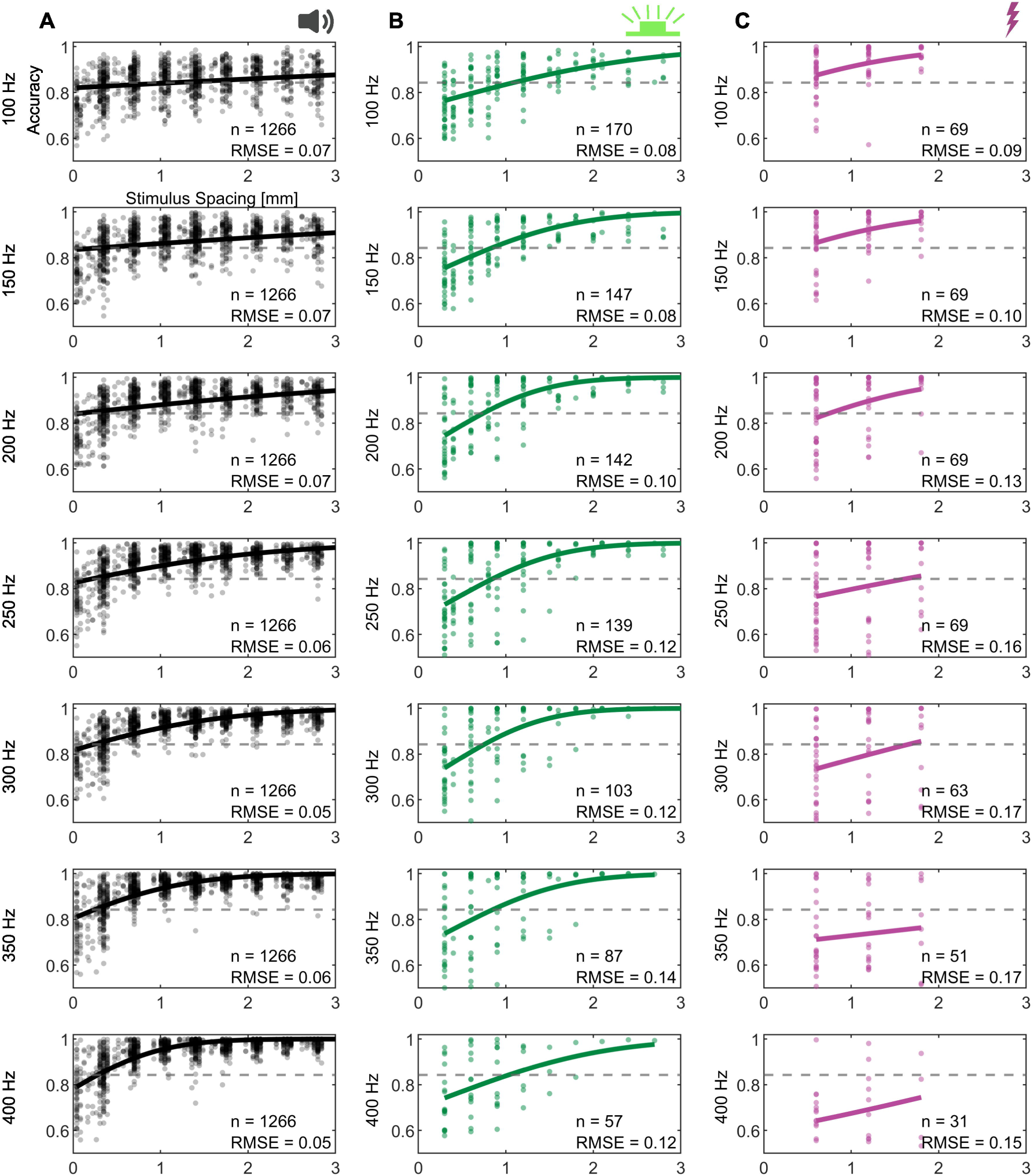
Neurometric functions at different activity levels for acoustic, electrical and optical stimulation. **(A–C)** Neurometric functions, as described in Fig. 8B, shown for different activity levels (rows) and stimulus modalities (columns, A–C). Background points show decoding accuracy for individual stimulus pairs at each FR, given next to each ordinate. The root mean square error (RMSE) of each fit and the number of data points are indicated in each panel. Icons were created in BioRender (see **Fig. 2**).

## References

Alekseev, A., Hunniford, V., Zerche, M., Jeschke, M., El May, F., Vavakou, A., Siegenthaler, D., Hüser, M.A., Kiehn, S.M., Garrido-Charles, A., Meyer, A., Rambousky, A., Alvanos, T., Witzke, I., Rojas-Garcia, K.D., Draband, M.D., Cyganek, L., Klein, E., Ruther, P., Huet, A., Trenholm, S., Macé, E., Kusch, K., Bruegmann, T., Wolf, B.J., Mager, T., Moser, T., 2025. Efficient and sustained optogenetic control of sensory and cardiac systems. Nat. Biomed. Eng 10, 277–292. 10.1038/s41551-025-01461-1

Auerbach, B.D., Radziwon, K., Salvi, R., 2019. Testing the Central Gain Model: Loudness Growth Correlates with Central Auditory Gain Enhancement in a Rodent Model of Hyperacusis. Neuroscience 407, 93–107. 10.1016/j.neuroscience.2018.09.036

Azees, A.A., Thompson, A.C., Thomas, R., Zhou, J., Ruther, P., Wise, A.K., Ajay, E.A., Garrett, D.J., Quigley, A., Fallon, J.B., Richardson, R.T., 2023. Spread of activation and interaction between channels with multi-channel optogenetic stimulation in the mouse cochlea. Hearing Research 440, 108911. 10.1016/j.heares.2023.108911

Bali, B., Gruber-Dujardin, E., Kusch, K., Rankovic, V., Moser, T., 2022. Analyzing efficacy, stability, and safety of AAV-mediated optogenetic hearing restoration in mice. Life Sci Alliance 5, e202101338. 10.26508/lsa.202101338

Bartels, M., Hernandez, V.H., Krenkel, M., Moser, T., Salditt, T., 2013. Phase contrast tomography of the mouse cochlea at microfocus x-ray sources. Appl Phys Lett 103, 083703. 10.1063/1.4818737

Bierer, J.A., 2010. Probing the electrode-neuron interface with focused cochlear implant stimulation. Trends Amplif 14, 84–95. 10.1177/1084713810375249

Bierer, J.A., 2007. Threshold and channel interaction in cochlear implant users: Evaluation of the tripolar electrode configurationa). J. Acoust. Soc. Am. 121.

Bonham, B.H., Litvak, L.M., 2008. Current focusing and steering: Modeling, physiology, and psychophysics. Hearing Research 242, 141–153. 10.1016/j.heares.2008.03.006

Dieter, A., Duque-Afonso, C.J., Rankovic, V., Jeschke, M., Moser, T., 2019. Near physiological spectral selectivity of cochlear optogenetics. Nat Commun 10, 1962. 10.1038/s41467-019-09980-7

Dieter, A., Klein, E., Keppeler, D., Jablonski, L., Harczos, T., Hoch, G., Rankovic, V., Paul, O., Jeschke, M., Ruther, P., Moser, T., 2020. μLED-based optical cochlear implants for spectrally selective activation of the auditory nerve. EMBO Mol Med 12, e12387. 10.15252/emmm.202012387

Drotos, A.C., Roberts, M.T., 2024. Identifying neuron types and circuit mechanisms in the auditory midbrain. Hear Res 442, 108938. 10.1016/j.heares.2023.108938

Fiáth, R., Márton, A.L., Mátyás, F., Pinke, D., Márton, G., Tóth, K., Ulbert, I., 2019. Slow insertion of silicon probes improves the quality of acute neuronal recordings. Sci Rep 9, 111. 10.1038/s41598-018-36816-z

Govorunova, E.G., Gou, Y., Sineshchekov, O.A., Li, H., Lu, X., Wang, Y., Brown, L.S., St-Pierre, F., Xue, M., Spudich, J.L., 2022. Kalium channelrhodopsins are natural light-gated potassium channels that mediate optogenetic inhibition. Nat Neurosci 25, 967–974. 10.1038/s41593-022-01094-6

Gradinaru, V., Thompson, K.R., Zhang, F., Mogri, M., Kay, K., Schneider, M.B., Deisseroth, K., 2007. Targeting and readout strategies for fast optical neural control in vitro and in vivo. The Journal of Neuroscience 27, 14231.

Harris, D.M., Shannon, R.V., Snyder, R., Carney, E., 1997. Multi-unit mapping of acoustic stimuli in gerbil inferior colliculus. Hearing Research 108, 145–156. 10.1016/S0378-5955(97)00047-6

Hernandez, V.H., Gehrt, A., Reuter, K., Jing, Z., Jeschke, M., Mendoza Schulz, A., Hoch, G., Bartels, M., Vogt, G., Garnham, C.W., Yawo, H., Fukazawa, Y., Augustine, G.J., Bamberg, E., Kügler, S., Salditt, T., De Hoz, L., Strenzke, N., Moser, T., 2014. Optogenetic stimulation of the auditory pathway. J Clin Invest 124, 1114–1129. 10.1172/JCI69050

Huet, A.T., Dombrowski, T., Rankovic, V., Thirumalai, A., Moser, T., 2021a. Developing Fast, Red-Light Optogenetic Stimulation of Spiral Ganglion Neurons for Future Optical Cochlear Implants. Front Mol Neurosci 14, 635897. 10.3389/fnmol.2021.635897

Huet, A.T., Dombrowski, T., Rankovic, V., Thirumalai, A., Moser, T., 2021b. Developing Fast, Red-Light Optogenetic Stimulation of Spiral Ganglion Neurons for Future Optical Cochlear Implants. Front. Mol. Neurosci. 14. 10.3389/fnmol.2021.635897

Huet, A.T., Mager, T., Gossler, C., Moser, T., 2024. Toward Optogenetic Hearing Restoration. Annual Review of Neuroscience. 10.1146/annurev-neuro-070623-103247

Huet, A.T., Rankovic, V., 2021. Application of Targeting-Optimized Chronos for Stimulation of the Auditory Pathway, in: Methods in Molecular Biology (Clifton, N.J.). pp. 261–285.

Hunniford, V., Kühler, R., Wolf, B., Keppeler, D., Strenzke, N., Moser, T., 2023. Patient perspectives on the need for improved hearing rehabilitation: A qualitative survey study of German cochlear implant users. Front Neurosci 17, 1105562. 10.3389/fnins.2023.1105562

Izzo, A.D., Richter, C.-P., Jansen, E.D., Walsh Jr., J.T., 2006. Laser stimulation of the auditory nerve. Lasers in Surgery and Medicine 38, 745–753. 10.1002/lsm.20358

Jablonski, L., Harczos, T., Wolf, B.J., Hoch, G., Khurana, L., Dieter, A., Roos, L., Hessler, R., Ayub, S., Ruther, P., Moser, T., 2025. Hearing restoration by a low-weight power-efficient multichannel optogenetic cochlear implant system. J. Neural Eng. 22. 10.1088/1741-2552/adf00f

Keppeler, D., Kampshoff, C.A., Thirumalai, A., Duque-Afonso, C.J., Schaeper, J.J., Quilitz, T., Töpperwien, M., Vogl, C., Hessler, R., Meyer, A., Salditt, T., Moser, T., 2021. Multiscale photonic imaging of the native and implanted cochlea. Proc Natl Acad Sci U S A 118, e2014472118. 10.1073/pnas.2014472118

Keppeler, D., Merino, R.M., Lopez de la Morena, D., Bali, B., Huet, A.T., Gehrt, A., Wrobel, C., Subramanian, S., Dombrowski, T., Wolf, F., Rankovic, V., Neef, A., Moser, T., 2018. Ultrafast optogenetic stimulation of the auditory pathway by targeting-optimized Chronos. EMBO J 37, e99649. 10.15252/embj.201899649

Keppeler, D., Schwaerzle, M., Harczos, T., Jablonski, L., Dieter, A., Wolf, B., Ayub, S., Vogl, C., Wrobel, C., Hoch, G., Abdellatif, K., Jeschke, M., Rankovic, V., Paul, O., Ruther, P., Moser, T., 2020. Multichannel optogenetic stimulation of the auditory pathway using microfabricated LED cochlear implants in rodents. Sci Transl Med 12, eabb8086. 10.1126/scitranslmed.abb8086

Khurana, L., Keppeler, D., Jablonski, L., Moser, T., 2022. Model-based prediction of optogenetic sound encoding in the human cochlea by future optical cochlear implants. Computational and Structural Biotechnology Journal 20, 3621–3629. 10.1016/j.csbj.2022.06.061

Kleinlogel, S., Feldbauer, K., Dempski, R.E., Fotis, H., Wood, P.G., Bamann, C., Bamberg, E., 2011. Ultra light-sensitive and fast neuronal activation with the Ca2+-permeable channelrhodopsin CatCh. Nat Neurosci 14, 513–518. 10.1038/nn.2776

Koert, E., Götz, J., Albrecht, N., Vavakou, A., Wolf, B.J., Moser, T., 2026. Optogenetic cochlear stimulation evokes midbrain activity with near-physiological temporal fidelity. 10.64898/2026.05.16.724905

Kral, A., Hartmann, R., Mortazavi, D., Klinke, R., 1998. Spatial resolution of cochlear implants: the electrical field and excitation of auditory afferents. Hear. Res. 121, 11–28. 10.1016/S0378-5955(98)00061-6

Kriegeskorte, N., 2008. Representational similarity analysis – connecting the branches of systems neuroscience. Front. Sys. Neurosci. 10.3389/neuro.06.004.2008

Lv, J., Wang, H., Cheng, X., Chen, Y., Wang, D., Zhang, L., Cao, Q., Tang, H., Hu, S., Gao, K., Xun, M., Wang, J., Wang, Z., Zhu, B., Cui, C., Gao, Z., Guo, L., Yu, S., Jiang, L., Yin, Y., Zhang, J., Chen, B., Wang, W., Chai, R., Chen, Z.-Y., Li, H., Shu, Y., 2024. AAV1-hOTOF gene therapy for autosomal recessive deafness 9: a single-arm trial. The Lancet 403, 2317–2325. 10.1016/S0140-6736(23)02874-X

Mager, T., Lopez de la Morena, D., Senn, V., Schlotte, J., D Errico, A., Feldbauer, K., Wrobel, C., Jung, S., Bodensiek, K., Rankovic, V., Browne, L., Huet, A., Jüttner, J., Wood, P.G., Letzkus, J.J., Moser, T., Bamberg, E., 2018. High frequency neural spiking and auditory signaling by ultrafast red-shifted optogenetics. Nat Commun 9, 1750. 10.1038/s41467-018-04146-3

Matic, A.I., Walsh, J.T., Jr, Richter, C.-P., 2011. Spatial extent of cochlear infrared neural stimulation determined by tone-on-light masking. J Biomed Opt 16, 118002. 10.1117/1.3655590

McInturff, S., Coen, F.-V., Hight, A.E., Tarabichi, O., Kanumuri, V.V., Vachicouras, N., Lacour, S.P., Lee, D.J., Brown, M.C., 2022. Comparison of Responses to DCN vs. VCN Stimulation in a Mouse Model of the Auditory Brainstem Implant (ABI). J Assoc Res Otolaryngol 23, 391–412. 10.1007/s10162-022-00840-8

Michael, M., Wolf, B.J., Klinge-Strahl, A., Jeschke, M., Moser, T., Dieter, A., 2023. Devising a framework of optogenetic coding in the auditory pathway: Insights from auditory midbrain recordings. Brain Stimul 16, 1486–1500. 10.1016/j.brs.2023.09.018

Micheyl, C., Xiao, L., Oxenham, A.J., 2012. Characterizing the dependence of pure-tone frequency difference limens on frequency, duration, and level. Hearing Research 292, 1–13. 10.1016/j.heares.2012.07.004

Middlebrooks, J.C., Snyder, R.L., 2007. Auditory prosthesis with a penetrating nerve array. J. Assoc. Res. Otolaryngol. 8, 258–279. 10.1007/s10162-007-0070-2

Miller, C.A., Abbas, P.J., Robinson, B.K., Nourski, K.V., Zhang, F., Jeng, F.-C., 2006. Electrical excitation of the acoustically sensitive auditory nerve: single-fiber responses to electric pulse trains. J Assoc Res Otolaryngol 7, 195–210. 10.1007/s10162-006-0036-9

Mittring, A., Moser, T., Huet, A.T., 2023. Graded optogenetic activation of the auditory pathway for hearing restoration. Brain Stimul 16, 466–483. 10.1016/j.brs.2023.01.1671

Moser, T., Chen, H., Kusch, K., Behr, R., Vona, B., 2024. Gene therapy for deafness: are we there now? EMBO Mol Med 16, 675–677. 10.1038/s44321-024-00058-6

Müller, M., 1996. The cochlear place-frequency map of the adult and developing Mongolian gerbil. Hear. Res 94, 148–156. 10.1016/0378-5955(95)00230-8

Richter, C.-P., Rajguru, S.M., Matic, A.I., Moreno, E.L., Fishman, A.J., Robinson, A.M., Suh, E., Walsh, J.T., 2011. Spread of cochlear excitation during stimulation with pulsed infrared radiation: inferior colliculus measurements. J Neural Eng 8, 056006. 10.1088/1741-2560/8/5/056006

Roos, L., Garrido-Charles, A., Albrecht, N., Vavakou, A., Alekseev, A., Bleyer, M., Thirumalai, A., Mittring, A., Alvanos, T., Huet, A.T., Bamberg, E., Kusch, K., Wolf, B.J., Moser, T., Mager, T., 2026. Channelrhodopsin variants for high-rate optogenetic neurostimulation at low light intensities. EMBO Mol Med 18, 462–491. 10.1038/s44321-025-00350-z

Sabesan, S., Fragner, A., Bench, C., Drakopoulos, F., Lesica, N.A., 2023. Large-scale electrophysiology and deep learning reveal distorted neural signal dynamics after hearing loss. eLife 12, e85108. 10.7554/eLife.85108

Schaeper, J.J., Kampshoff, C.A., Wolf, B.J., Roos, L., Michanski, S., Ruhwedel, T., Eckermann, M., Meyer, A., Jeschke, M., Wichmann, C., Moser, T., Salditt, T., 2025. 3D virtual histology of rodent and primate cochleae with multi-scale phase-contrast X-ray tomography. Sci Rep 15, 7933. 10.1038/s41598-025-89431-0

Schaeper, J.J., Liberman, M.C., Salditt, T., 2023. Imaging of excised cochleae by micro-CT: staining, liquid embedding, and image modalities. JMI 10, 053501. 10.1117/1.JMI.10.5.053501

Schnupp, J.W.H., Garcia-Lazaro, J.A., Lesica, N.A., 2015. Periodotopy in the gerbil inferior colliculus: local clustering rather than a gradient map. Front Neural Circuits 9, 37. 10.3389/fncir.2015.00037

Shannon, R.V., 1983. Multichannel electrical stimulation of the auditory nerve in man. II. Channel interaction. Hear. Res. 12, 1–16.

Snyder, R.L., Bierer, J.A., Middlebrooks, J.C., 2004. Topographic spread of inferior colliculus activation in response to acoustic and intracochlear electric stimulation. J. Assoc. Res. Otolaryngol 5, 305–322. 10.1007/s10162-004-4026-5

Stringer, C., Pachitariu, M., 2022. Cellpose 2.0: how to train your own model. bioRxiv 2022.04.01.486764. 10.1101/2022.04.01.486764

Stringer, C., Wang, T., Michaelos, M., Pachitariu, M., 2021. Cellpose: a generalist algorithm for cellular segmentation. Nat Methods 18, 100–106. 10.1038/s41592-020-01018-x

Thirumalai, A., Henseler, J., Enayati, M., Kusch, K., Hessler, R., Moser, T., Huet, A.T., 2025. Improved optogenetic modification of spiral ganglion neurons for future optical cochlear implants. Theranostics 15, 4270–4286. 10.7150/thno.104474

Vollmer, M., Beitel, R.E., Snyder, R.L., Leake, P.A., 2007. Spatial selectivity to intracochlear electrical stimulation in the inferior colliculus is degraded after long-term deafness in cats. J Neurophysiol 98, 2588–2603. 10.1152/jn.00011.2007

WHO, 2021. Deafness and hearing loss. WHO.

Wier, C.C., Jesteadt, W., Green, D.M., 1977. Frequency discrimination as a function of frequency and sensation level. J. Acoust. Soc. Am. 61, 178–184. 10.1121/1.381251

Wohlbauer, D.M., Dillier, N., 2025. A Hundred Ways to Encode Sound Signals for Cochlear Implants. Annual Review of Biomedical Engineering. 10.1146/annurev-bioeng-102623-121249

Wrobel, C., Dieter, A., Huet, A., Keppeler, D., Duque-Afonso, C.J., Vogl, C., Hoch, G., Jeschke, M., Moser, T., 2018. Optogenetic stimulation of cochlear neurons activates the auditory pathway and restores auditory-driven behavior in deaf adult gerbils. Sci Transl Med 10, eaao0540. 10.1126/scitranslmed.aao0540

Zeng, F.G., 2017. Challenges in Improving Cochlear Implant Performance and Accessibility. IEEE Trans Biomed Eng 64, 1662–1664. 10.1109/TBME.2017.2718939

Zhu, X., Huang, L., Zheng, Y., Song, Y., Xu, Q., Wang, J., Si, K., Duan, S., Gong, W., 2019. Ultrafast optical clearing method for three-dimensional imaging with cellular resolution. Proc Natl Acad Sci U S A 116, 11480–11489. 10.1073/pnas.1819583116

