## Supplementary Figures and Methods for "Multichannel optical cochlear implants enable spectrally distinct auditory activity"

**Albrecht et al.**

### Appendix Figures

**Appendix Figure S1. Placement of the recording array along the tonotopic axis of the ICC.** **page 2**

**Appendix Figure S2. Spontaneous ICC activity across stimulation-modality groups.** **page 2**

**Appendix Figure S3. Balanced vs. unbalanced averaging of the rate-level functions** **page 3**

**Appendix Figure S4. Effect of analysis method on dynamic-range estimates.** **page 3**

**Appendix Figure S5. Thresholds for ICC activation by pure tones.** **page 4**

**Appendix Figure S6. Comparison of thresholds for the detection of ICC activity in response to stimulation by individual LEDs along the tonotopic axis.** **page 5**

**Appendix Figure S7. Balanced vs. unbalanced analysis of spectral spread.** **page 5**

**Appendix Figure S8. Alternative estimation of spectral spread of cochlear excitation by cumulative d-Prime-Analysis as in Dieter et al., 2019.** **page 6**

**Appendix Figure S9. Estimation of an equivalent SPL by d-Prime-Analysis.** **page 7**

**Appendix Figure 10. Automated spectral-peak detection method.** **page 9**

### Appendix Methods

**Quantification of response strength and spectral spread using d'-Analysis** **page 10**

### Appendix Figures

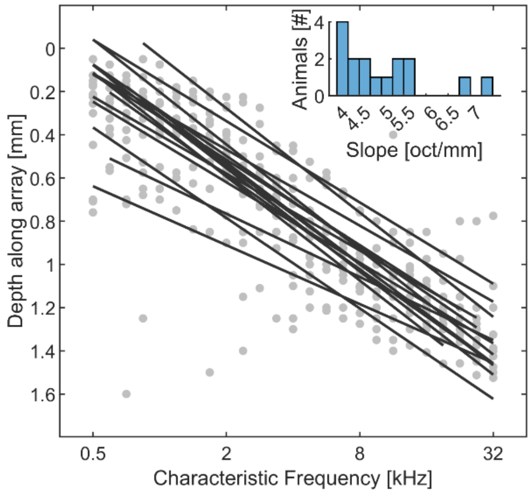


**Appendix Figure S1. Placement of the recording array along the tonotopic axis of the ICC.** Depth of the best electrode for the presented sound frequencies relative to the most dorsal recording electrode Tonotopic slopes were estimated by linear fits for each animal after outlier removal (N = 8 non-injected and N = 8 ChReef-injected animals). The inset axis shows the distribution of tonotopic slopes across animals.

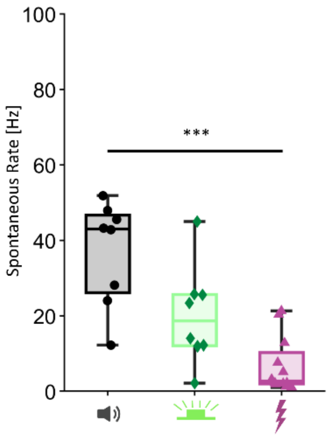


**Appendix Figure S2. Spontaneous ICC activity across stimulation-modality groups.** Spontaneous firing rates recorded in the ICC from ChReef-injected animals used for optical stimulation (*N* = 8), non-injected animals used for acoustic stimulation (*N* = 8), and non-injected animals used for electrical stimulation (N = 12). Multiple frequencies or emitters were tested within individual animals. Each data point represents the average across recordings in one animal. Groups were compared after averaging recordings obtained from the same animal using a Kruskal–Wallis test followed by Tukey's multiple-comparisons test. Box plots indicate quartiles and medians; whiskers extend to minimum and maximum values. Only significant differences are indicated (*p < 0.05, **p < 0.01, ***p < 0.001). Exact p-values are reported in **Table EV1**. Throughout the figure, acoustic data are shown in black with round markers, optical data in green with diamond markers, and electrical data in purple with triangle markers. Icons were created in BioRender (see. **Fig. 2**).

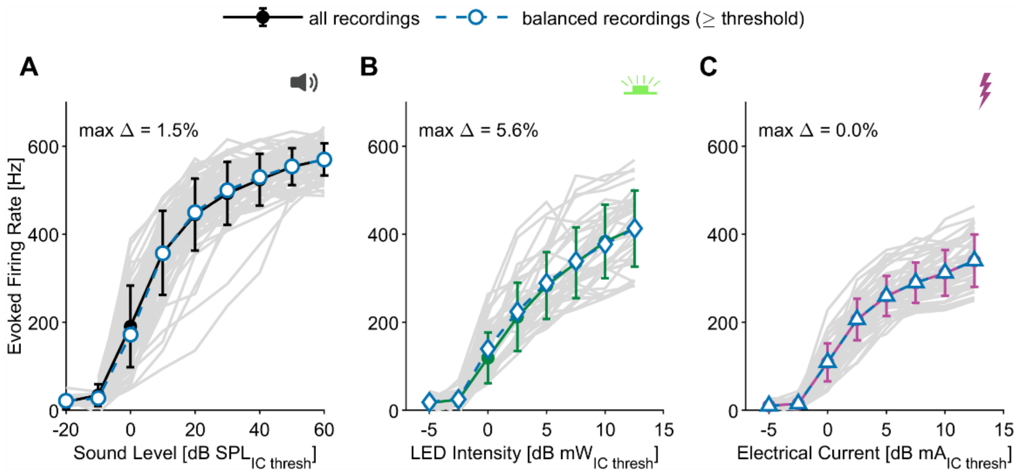


**Appendix Figure S3. Balanced vs. unbalanced averaging of the rate-level functions.** Mean onset-evoked firing rate (± SD) vs. intensity relative to ICC threshold for (**A**) acoustic, (**B**) LED, and (**C**) electric (eCI) stimulation. "All recordings" (colored) uses every recording available in each bin , whereas "balanced recordings" (blue dashed) includes only recordings with data available in every plotted suprathreshold bin. Grey lines indicate individual recordings included in the balanced subset. Max Δ, maximum difference between the two averages across supra-threshold bins. Number of recordings: acoustic, total n = 236, balanced n = 73 from 8 gerbils; LED, total n = 59, balanced n = 34 from 8 gerbils; electrical, total n = 47, balanced n = 47 from 12 gerbils.

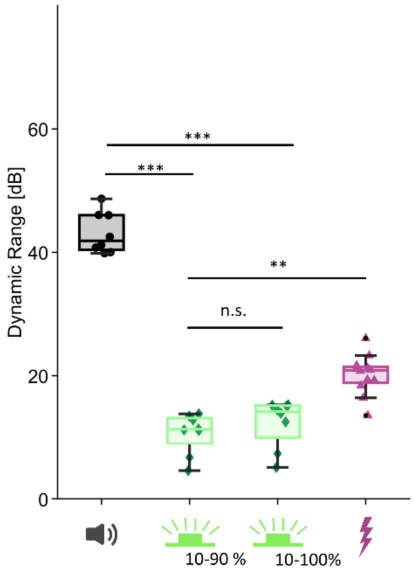


**Appendix Figure S4. Effect of analysis method on dynamic-range estimates.**

Distribution of dynamic ranges for acoustic, optical and electrical stimulation. For optical stimulation, the dynamic range was additionally estimated as the intensity range required to increase firing rates from 10% to 100% of the maximum firing rate. Multiple frequencies or emitters were tested within individual animals (N = 8/8/12 for acoustic/optical/electrical stimulation). Each data point represents the average across recordings in one animal. Groups were compared after averaging recordings obtained from the same animal using a Kruskal–Wallis test followed by Tukey's multiple-comparisons test. Only significant differences are indicated (*p < 0.05, **p < 0.01, ***p < 0.001). Exact p-values are reported in **Table EV1**. Box plots indicate quartiles and medians; whiskers extend to minimum and maximum values. Throughout the figure, acoustic data are shown in black with round markers, optical data in green with diamond markers, and electrical data in purple with triangle markers. Icons were created in BioRender (see. **Fig. 2**).

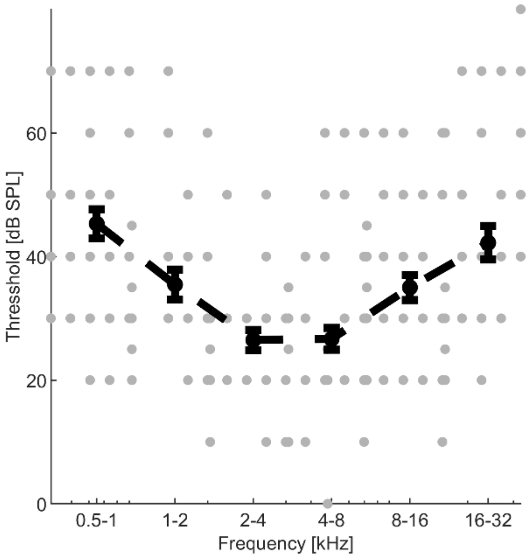


**Appendix Figure S5. Thresholds for ICC activation by pure tones.**

Threshold of ICC activation by pure tones of different frequencies in non-injected animals (N = 8), defined as the first stimulation intensity to elicit a d'≥1. Values were binned by frequency at a width of one octave (mean ± SEM).

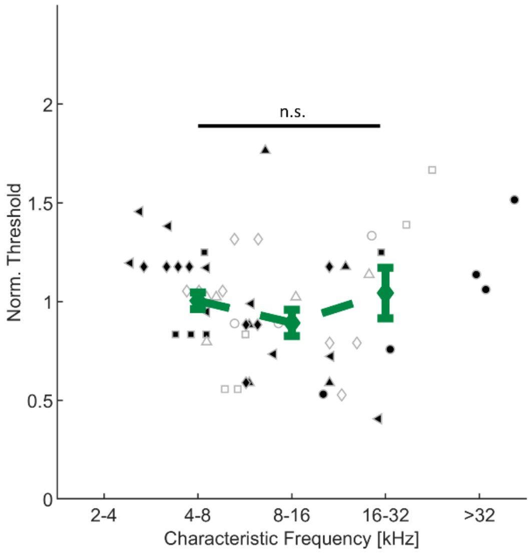


**Appendix Figure S6. Comparison of thresholds for the detection of ICC activity in response to stimulation by individual LEDs along the tonotopic axis.**

A $C_f$ for each LED was determined from the centroid of the corresponding STC and thresholds were normalized to the average threshold in each animal (**Fig. 1C**). Groups corresponding to octave bands (4–8, 8–16, and 16–32 kHz) were compared using a Kruskal–Wallis's test followed by Tukey's post hoc comparisons (n.s., non-significant). Data from 59 LED recordings in 8 gerbils are shown. Exact p-values are reported in **Table EV1**.

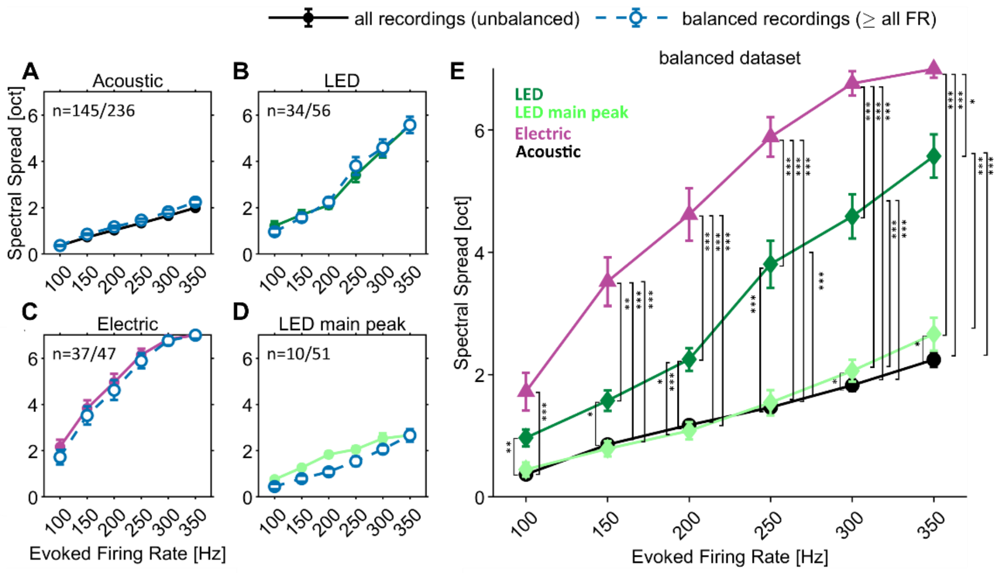


**Appendix Fig. S7. Balanced vs. unbalanced analysis of spectral spread.**

(**A-D**) Mean spectral spread (± SEM, octaves) as a function of activity level (evoked firing rate

[Hz]) for (**A**) acoustic, (**B**) LED, (**C**) electric and (**D**) the LED main spectral peak. "All recordings" (colored) averages every recording available at each firing rate (unbalanced); "balanced" (blue dashed) averages only the recordings that provide a spectral spread value at every firing rate (n = balanced/total per panel).

(**E**) Comparative overview of the balanced dataset for all modalities. Statistical comparisons were conducted using a linear mixed-effects model, with animal treated as a random factor and evoked firing rate and modality treated as categorical fixed effects as in **Fig. 6**, (*p<0.05, **p<0.01, **p<0.001). Exact p-values are reported in **Table EV1**. Evoked firing rate was modeled as a categorical factor to avoid assuming a linear relationship between firing rate and the outcome. Only significant effects are indicated. The number of recordings contributing to each activity level is shown in **A-D** and was obtained from N = 8/8/12 gerbils for acoustic/optical/electrical stimulation, respectively.

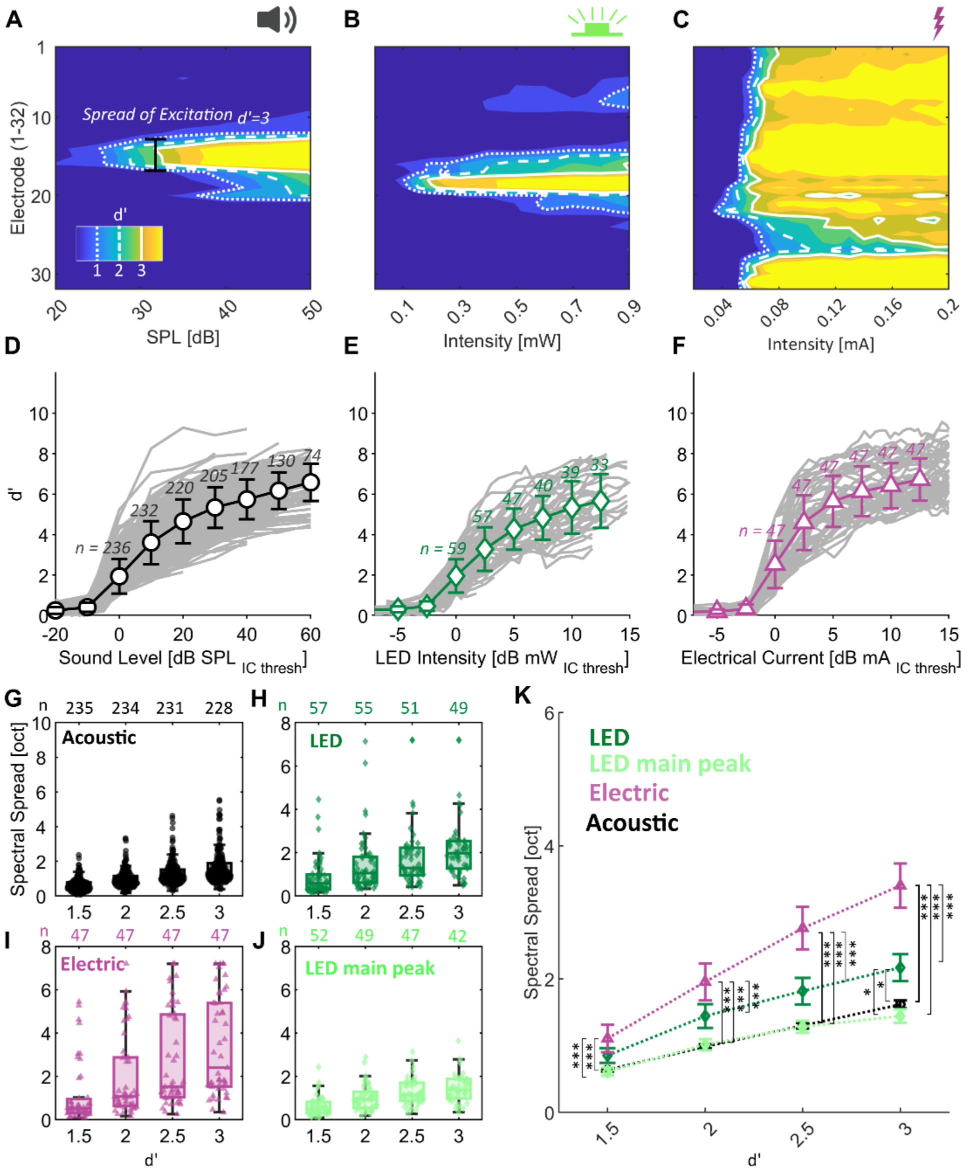

**Appendix Figure S8. Alternative estimation of spectral spread of cochlear excitation by cumulative d-Prime-Analysis as in Dieter et al., 2019.**

**(A-C**) D-Prime-based STCs for sound, light and electrical stimulation. Quantification of spectral spread in different stimulation modalities: Spread of excitation (SoE) is measured as the width of the STC at different activation levels quantified as the highest cumulative d-Prime in response to a given stimulus (example shown for d'=3 in Panel A). Icons were created in BioRender (see. **Fig. 2**).

**(D-F)** Cumulative d-Prime quantified using the maximally responding electrode at each stimulus intensity (mean ± SD) for acoustic (**D**), optical (**E**), and electrical (**F**) stimulation. Traces for individual recordings are plotted in the background. Because multiple frequencies or emitters were tested within individual animals, the number above each error bar indicates the number of recordings contributing to that data point.

**(G-J)** Spectral Spread at different d-Prime-Levels for acoustic (**D**), optical (**E**) and electrical (**F**) stimulation. For optical stimulation the spectral spread was additionally quantified while excluding any secondary spectral peaks. Because multiple frequencies or emitters were tested within individual animals, the number above each error bar indicates the number of recordings contributing to that data point.

**(K)** Comparison of spectral spread (Mean ± SEM) for different stimulus modalities i.e. sound, light and electrical stimulation as shown in **D-G**. To facilitate comparison, the d'-levels for comparison as well as the statistical analysis were matched to Dieter et al., 2019**.** Thus, statistical comparisons were conducted by repeated-measures ANOVA followed by Tukey's post-hoc comparison. (*p<0.05, **p<0.01, **p<0.001). Only significant differences are indicated. Data were acquired from N = 8/8/12 gerbils for acoustic/optical/electric stimulation, respectively. Exact p-values are reported in **Table EV1**. Throughout the figure, acoustic data are shown in black with round markers, optical data in green with diamond markers, and electrical data in purple with triangle markers.

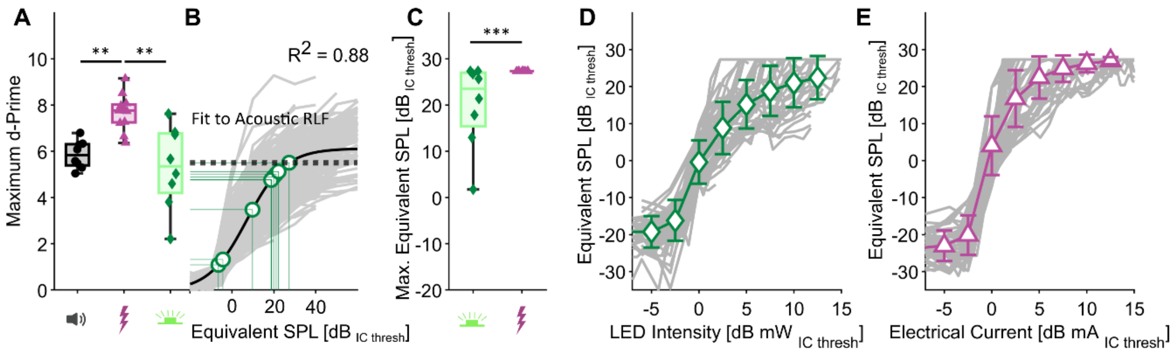


**Appendix Figure S9. Estimation of an equivalent SPL by d-Prime-Analysis.**

**(A)** Distribution of the maximum cumulative d-Prime achieved by acoustic, optical, and electrical stimulation.

**(B)** Quantification of an equivalent sound pressure level (SPL) for optical and electrical stimulation. A sigmoid fit to the dPrime-level function for acoustic stimulation (**Fig. S9**) (pseudo-$R^2$ =0.88) was used as a reference to map firing rates elicited by optical and electrical stimulation onto an equivalent SPL.

**(C)** Comparison of the maximum equivalent SPL, derived from the mapping in panel **B**, achieved by optical and electrical stimulation.

**(D-E)** Mapping of evoked firing rates achieved by optical (**J**) and electrical stimulation (**K**) to an equivalent SPL (mean ± SD). The data shown in **Appendix Fig. S8D-F** was mapped onto the dPrime-level function for acoustic stimulation as shown in B. Traces for individual recordings are shown in the background. The number of recordings contributing to each data point corresponds to the values shown in Fig. S8D

Statistical significance is indicated by stars (*p < 0.05, **p < 0.01, ***p < 0.001), based on Kruskal-Wallis test with Tukey's post-hoc pairwise comparison after averaging recordings obtained from the same animal. Exact p-values are reported in **Table EV1**. Boxes in **A** and **C** indicate quartiles and median, with whiskers extending to the minimum and maximum values. Multiple frequencies or emitters were tested within individual animals. Each data point in **A** and **C** represents the average across recordings in one animal. Throughout the figure, acoustic data are shown in black with round markers, optical data in green with diamond markers, and electrical data in purple with triangle markers. Icons in **A** and **C** were created in BioRender (see. **Fig. 2**).

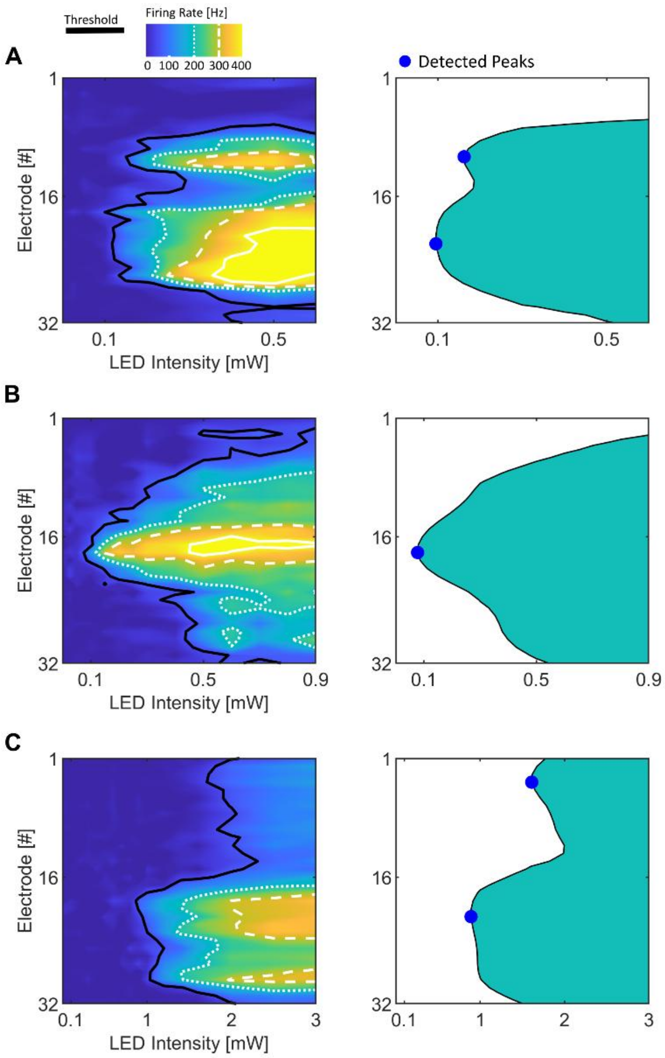


**Appendix Figure S10. Automated spectral-peak detection method. (A–C)** Threshold (d’ ≥ 1) contours of STCs obtained with single-LED stimulation are shown before smoothing in the left column and after smoothing with a two-dimensional Gaussian window in the right column. Smoothing was applied to reduce over-detection of noise-related peaks. Remaining peaks were automatically detected using MATLAB’s *findpeaks* function and are indicated by blue dots.

### Appendix Methods

#### Quantification of response strength and spectral spread using d'-Analysis

To enable comparison with previous studies, we additionally quantified ICC responses using cumulative d′ analysis, a measure derived from signal detection theory (Macmillan and Creelman, 2004). For the main figures, however, we used evoked firing rate to quantify response strength, because this allowed direct comparison to rate-level functions across modalities and avoided potential bias arising from differences in inter-trial variability between stimulus conditions.

Responses obtained from 30 repetitions at each stimulus intensity were used to calculate cumulative d′ values (Macmillan and Creelman, 2004; Middlebrooks and Snyder, 2007). For each recording site, firing-rate distributions evoked at successive stimulus intensities were used to construct empirical receiver operating characteristic (ROC) curves, from which the corresponding areas under the curve (AUC) were obtained. d′ values were then derived from the AUC using the inverse cumulative normal distribution and summed cumulatively across increasing stimulus intensities to yield the cumulative d′ value. The spatial distribution of cumulative d′ values across recording sites was visualized by drawing iso-contour lines at defined d′ levels (d′ = 1, 2, and 3) using MATLAB's built-in contourf function.

To compare the spread of excitation in this framework, we measured the width of the STC constructed from d'-contours at matched activation levels (Dieter et al., 2019; Keppeler et al., 2020). For each activation level, the corresponding stimulus intensity was defined as the lowest intensity at which a given d'-contour (d′ = 1.5 to 3) was reached. STC width was then measured as the distance between the most dorsal and ventral intersections of the cumulative d'=1-contour at that intensity. For this analysis, a cumulative d'-value of 1 served as the threshold criterion while the main analysis used a baseline d'-values (see Methods).
